# Cryo-EM structure of the Spo11 core complex bound to DNA

**DOI:** 10.1101/2023.10.31.564985

**Authors:** You Yu, Juncheng Wang, Kaixian Liu, Zhi Zheng, Meret Arter, Corentin Claeys Bouuaert, Stephen Pu, Dinshaw J. Patel, Scott Keeney

## Abstract

The DNA double-strand breaks that initiate meiotic recombination are formed by topoisomerase relative Spo11, supported by conserved auxiliary factors. Because high-resolution structural data are lacking, many questions remain about the architecture of Spo11 and its partners and how they engage with DNA. We report cryo-EM structures at up to 3.3 Å resolution of DNA-bound core complexes of *Saccharomyces cerevisiae* Spo11 with Rec102, Rec104, and Ski8. In these structures, monomeric core complexes make extensive contacts with the DNA backbone and with the recessed 3’-OH and first 5’ overhanging nucleotide, definitively establishing the molecular determinants of DNA end-binding specificity and providing insight into DNA cleavage preferences in vivo. The structures of individual subunits and their interfaces, supported by functional data in yeast, provide insight into the role of metal ions in DNA binding and uncover unexpected structural variation in homologs of the Top6BL component of the core complex.

## Introduction

Spo11 is related to the DNA-cleaving A subunit (Top6A) of archaeal topoisomerase VI ^1^. Both enzymes cut DNA through a transesterification reaction in which a tyrosine side chain severs the DNA backbone and attaches covalently to the 5′ terminus by a tyrosyl phosphodiester linkage. Two Top6A or Spo11 proteins work together to make a DSB with two-nucleotide 5′ overhangs ^2,3^.

Topo VI is a heterotetramer of two A and two B subunits. Top6A has two catalysis-critical domains—a winged-helix (WH) domain that carries the DNA cleaving tyrosine, and a metal-ion-binding Rossmann fold known as the Toprim domain ^4–6^ (**Fig. 1A**). Scission of each DNA strand involves the tyrosine of one Top6A monomer interacting with the Mg^2+^-binding pocket of the second Top6A monomer to form a hybrid active site, thus requiring dimer formation for catalysis. Sequence alignments and computational modeling by AlphaFold2 predict equivalent domains in Spo11 ^7–10^ (**Fig. 1A, B**).

**Fig. 1.**
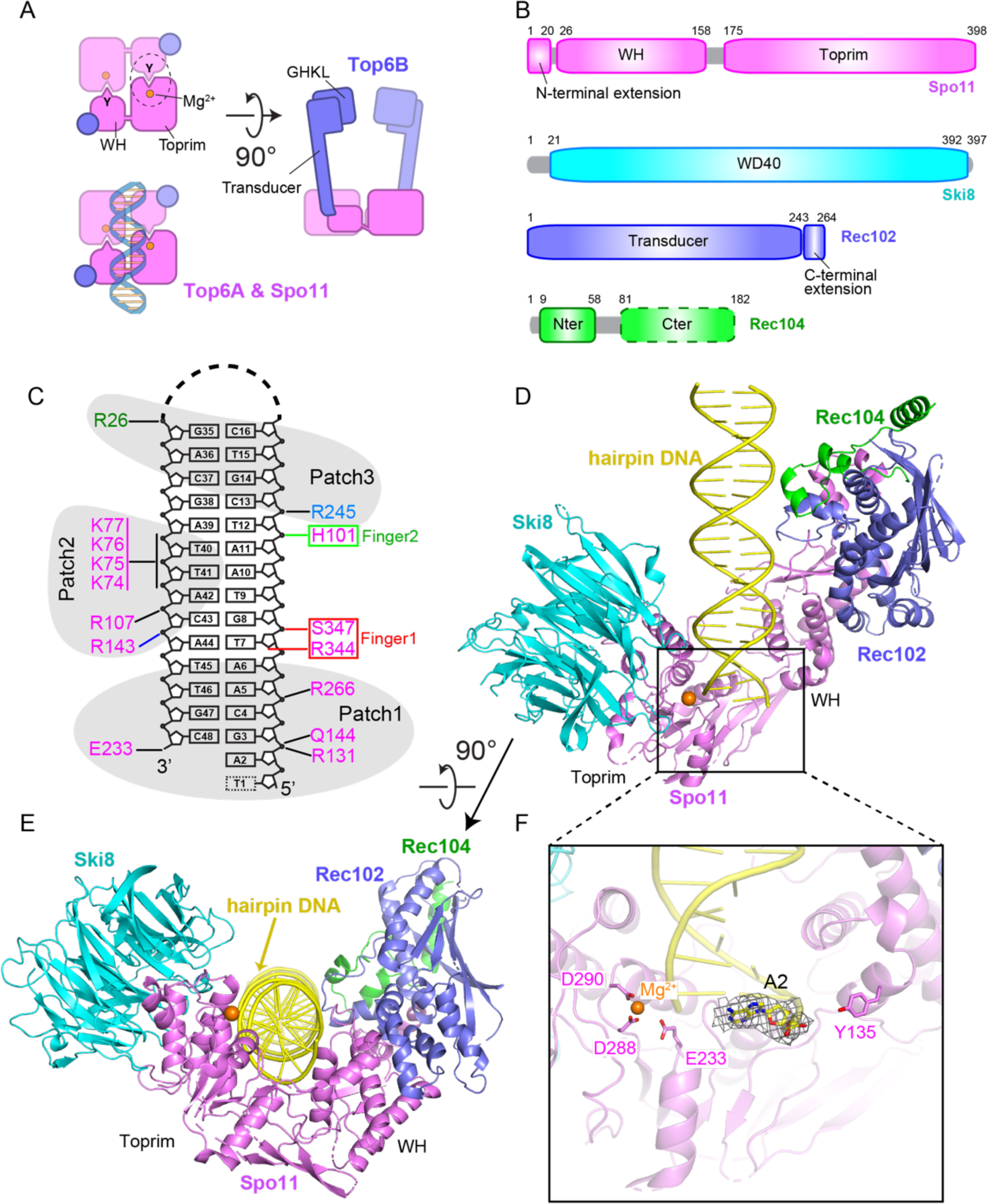
Cryo-EM structure of Spo11 core complex bound to hairpin DNA. (A) Schematic of tertiary and quaternary organization of the Topo VI holoenzyme. Left: top views (with and without DNA) looking down into the DNA-binding channel of Top6A (or Spo11), illustrating the dimer interface, catalytic tyrosine (Y), metal binding pocket, and hybrid active site (dashed circle). Right: side view. (B) Domain organization of Spo11, Ski8, Rec102 and Rec104. (C) Hairpin DNA sequence and intermolecular contacts between the DNA (mostly sugar-phosphate backbone) and amino acid side chains of Spo11 (magenta), Rec102 (blue), and Rec104 (green). Groups of DNA-interacting residues described in the text (patches and fingers) are highlighted (D and E) Two views of a ribbon representation of the 3.7 Å cryo-EM structure. (F) Detailed view of Spo11 and the 5’-overhang end of the hairpin DNA, highlighting select active site residues. Density of the first nucleotide of the overhang (A2) is shown as gray mesh.

Top6B has an N-terminal GHKL-family ATPase domain that dimerizes upon binding to ATP and a “transducer” domain consisting of a β sheet backed by three α helices, one of which provides a long lever arm connecting the GHKL domain to the Top6A binding interface (**Fig. 1A**) ^4,5,11^. Mouse and plant homologs of Top6B (Top6BL) contain domains predicted to resemble both the transducer and GHKL folds, but the homologs from some species, including yeasts, resemble only the transducer domain ^1,12,13^. Top6B also has a helix-2-turn-helix (H2TH) domain and a C-terminal domain (CTD); the function of these is unknown. The eukaryotic homologs lack the CTD and current homology models differ as to whether an H2TH domain is present.

In *S. cerevisiae*, Spo11 forms a “core complex” with Rec102, Rec104, and Ski8 ^14^. Spo11 and its partners are conserved in most eukaryotic taxa, but with substantial sequence diversity ^15–18^. Purified recombinant core complexes have a 1:1:1:1 stoichiometry, which was unexpected because Top6A forms a stable dimer ^4–6^ and because DNA cleavage requires hybrid active sites. Rec102 is homologous to the Top6B transducer domain, but lacks a clear equivalent of the GHKL domain ^1,16^ (**Fig. 1B**). Loss of the GHKL domain is an evolutionarily common scenario in eukaryotic homologs, having occurred independently in multiple lineages^16^.

Rec104 has been enigmatic: its apparently narrow phylogenetic distribution ^15^ and consequent shallow multiple sequence alignment have thus far precluded a high-confidence prediction by AlphaFold2. It has been hypothesized that Rec104 may have either evolved from or replaced the GHKL domain ^1,13,14,16^, but this remains to be verified.

Yeast Spo11 core complexes bind noncovalently with high affinity (sub-nanomolar *K_d_*) to DNA ends that mimic the cleavage product in having a two-nucleotide 5’ overhang ^14^. Blunt DNA ends or 3′ overhangs are bound with much lower affinity. This tight binding has been proposed to allow Spo11 to cap DSB ends, potentially affecting subsequent repair ^14,19,20^.

Crystal structures have been reported for Topo VI holoenzymes or individual subunits ^4–6,11^, but not for eukaryotic Spo11. There are homology models for yeast Spo11 core complexes templated on Topo VI ^12–14^ or predicted by RosettaFold and AlphaFold2 ^9^. However, these models differ from one another for subunit tertiary structure and protein-protein contacts ^9^, and no high-confidence model is yet available for Rec104. Moreover, no structure determination has been reported for any Spo11 or Topo VI relative bound to DNA, so the basis of DNA substrate selectivity is unknown. We address these issues by reporting cryo-EM structures of DNA-bound yeast Spo11 core complexes.

## Results and Discussion

### Cryo-EM structure of the Spo11 core complex bound to hairpin DNA

*S. cerevisiae* core complexes by themselves were conformationally variable in negative-stain EM experiments, but they adopted a more uniform, compact structure when bound to DNA^14^. Therefore, we leveraged the tight binding to DNA ends to determine a cryo-EM structure.

Core complexes were purified after expression in insect cells (**Fig. S1A**) and incubated in the presence of Mg^2+^ with a 23-bp hairpin DNA that had a two-nucleotide 5′ overhang (**Fig. 1C and Fig. S1B**). The mixture was loaded on cryo-EM grids without further purification. The structure of the core complex with bound hairpin DNA was solved at 3.7 Å resolution (cryo-EM reconstruction in **Fig. S2A–D**; statistics in **Table S1**; density representation in **Fig. S2E**). The model was built based on subunit structures predicted by AlphaFold2 ^10^, and manually refined by positioning of bulky amino acid side chains.

The core complex is “monomeric” in the structure (1:1:1:1 stoichiometry) and adopts a V-shaped topology shown in two orientations in a ribbon representation (**Fig. 1D, E**). Spo11 is located at the base of the complex and forms contacts with the three other subunits. The overhang end of the hairpin DNA is anchored in a deep cleft positioned between the WH and Toprim domains of Spo11 (**Fig. 1D**). The remainder of the duplex makes non-sequence-specific contacts primarily with Spo11 and fewer with both Rec102 and Rec104, while no interaction is observed between the DNA and Ski8 (**Fig. 1E**). The proteins form a left-handed wrap around the DNA. Protein-DNA contacts in the complex are summarized in **Fig. 1C** and discussed further below.

We could unambiguously trace the base, sugar and 5′-phosphate of the first nucleotide (A2) in the overhang adjacent to the terminal base pair. By contrast, the next overhang base (T1) could not be traced because of weak density, most likely due to flexibility. The catalytic residues — tyrosine Y135 involved in DNA covalent bond formation on the one hand, and the Mg^2+^-coordinating E233, D288 and D290 residues on the other — are positioned on opposite sides of the overhang end of the hairpin (**Fig. 1F**). This separation of active site components is as expected given that, with only one copy of Spo11, the structure shows the configuration of two catalytic half-sites (**Fig. 1A**). We consider it likely that the observed structure mimics the post-DSB product complex.

### Cryo-EM structure of the core complex bound to gapped DNA

We also solved the cryo-EM structure of the core complex bound in the presence of Mg^2+^ to a double-hairpin substrate with a ssDNA gap (**Fig. 2A and S1C**) at an improved resolution of 3.3 Å (cryo-EM reconstruction in **Fig. S3**; statistics in **Table S1**; and density maps in **Fig. S4A–E**). Two views are shown in a ribbon representation in **Fig. 2B, C**.

**Fig. 2.**
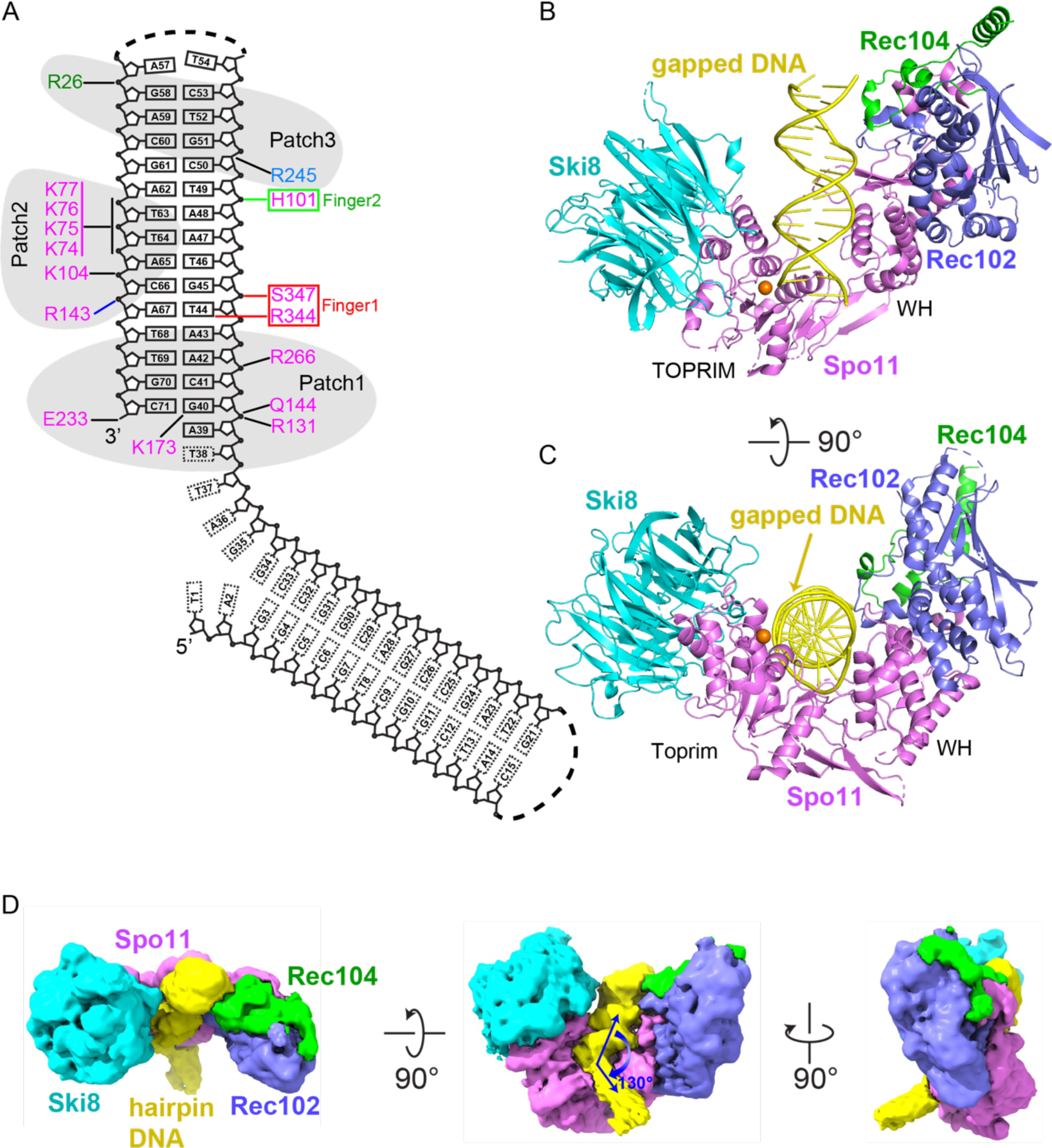
Cryo-EM structure Spo11 core complex bound to gapped DNA. (A) Gapped DNA sequence and intermolecular contacts between the DNA and amino acid side chains of Spo11 (magenta), Rec102 (blue), and Rec104 (green) (B and C) Two views of a ribbon representation of the 3.3 Å cryo-EM structure. (D) Three views of the unsharpened cryo-EM map.

We observed one hairpin bound to the core complex, allowing tracing of its duplex segment and the first nucleotide of the ssDNA (A39) including its 5′-phosphate. Notably, the bound hairpin is the one with a free 3′-OH end and ssDNA extending from the 5′ end (**Fig. 2A**), i.e., structurally analogous to the single-hairpin substrate initially used. We can trace weak density in the unsharpened map spanning the single-stranded nucleotides T38-T37-A36 and a portion of the duplex segment of the second hairpin, with the two duplex segments aligned at a relative angle of 130° (**Fig. 2D**). Protein-DNA contacts are summarized in **Fig. 2A**.

The distributions of α-helical and β-strand segments for the four subunits in the DNA-bound complex are shown in **Fig. S5**. Spo11 serves as a central hub through extensive protein-protein and protein-DNA contacts in the complex. The two arms of the V-shaped architecture of the protein scaffold comprise the WH domain of Spo11 interacting with Rec102 and Rec104, and the Toprim domain interacting with Ski8 (**Fig. 2B, C**). Because there is good superposition between the structures with hairpin DNA and gapped DNA (rmsd = 1.23 Å), the following sections on subunit architecture and protein-protein and protein-DNA interactions will focus on the higher resolution structure with gapped DNA.

### Validation of the Spo11–Ski8 interface

The interface of Spo11 with WD-repeat protein Ski8 was previously modeled using a crystal structure of the Ski complex, in which two copies of Ski8 contact two copies of a QRxxΦ motif in Ski3 that is also present in fungal Spo11 proteins ^14,21^. This motif is critical for Spo11– Ski8 interaction in *S. cerevisiae* both in vivo and in vitro ^14,22^. This same interface was also predicted computationally ^9^.

Consistent with these predictions, Ski8 in the cryo-EM structures adopts a propeller folding topology formed by seven WD40 repeats and recognizes the sequence Q_376_REIFF in the Spo11 Toprim domain (**Fig. S6A**). The interface is mediated by both hydrophobic and hydrogen bonding interactions. The hydrophobic core is formed by Spo11 residues F380 and F381 and surrounding Ski8 aromatic residues involved in CH-π and π-π stacking interactions (**Fig. S6B**). A hydrogen bonding network surrounds this hydrophobic core, including bonds between Q376 and R377 of Spo11 and Ski8 residues S63 and D18, respectively (**Fig. S6C**). Computational prediction had also suggested that an extended loop in Ski8 expands the interface with Spo11 ^9^. In agreement, we observed additional hydrogen bonding interactions that further stabilize the interface (**Fig. S6D**).

### Rec102 and its interaction with Spo11

Rec102 comprises a six-stranded β sheet extended by multiple α helices (**Fig. 2B, 3A, and S5C**). It has structural elements in common with the transducer domain of archaeal Top6B, as anticipated, but with substantial differences. We consider here five main features from Top6B: a central β sheet; a WKxY motif that is part of a proposed DNA binding surface; a “switch loop” that interacts with the ATP binding site in the GHKL domain; the long α-helical lever arm (“stalk”) that connects to Top6A; and the interface with Top6A (**Fig. 3A**). For this discussion, conserved β strands will be indicated by uppercase letters ordered by tertiary structure position; numerical designations for β strands will be according to primary structure position (e.g., **Fig. S5**), so these differ between species.

**Fig. 3.**
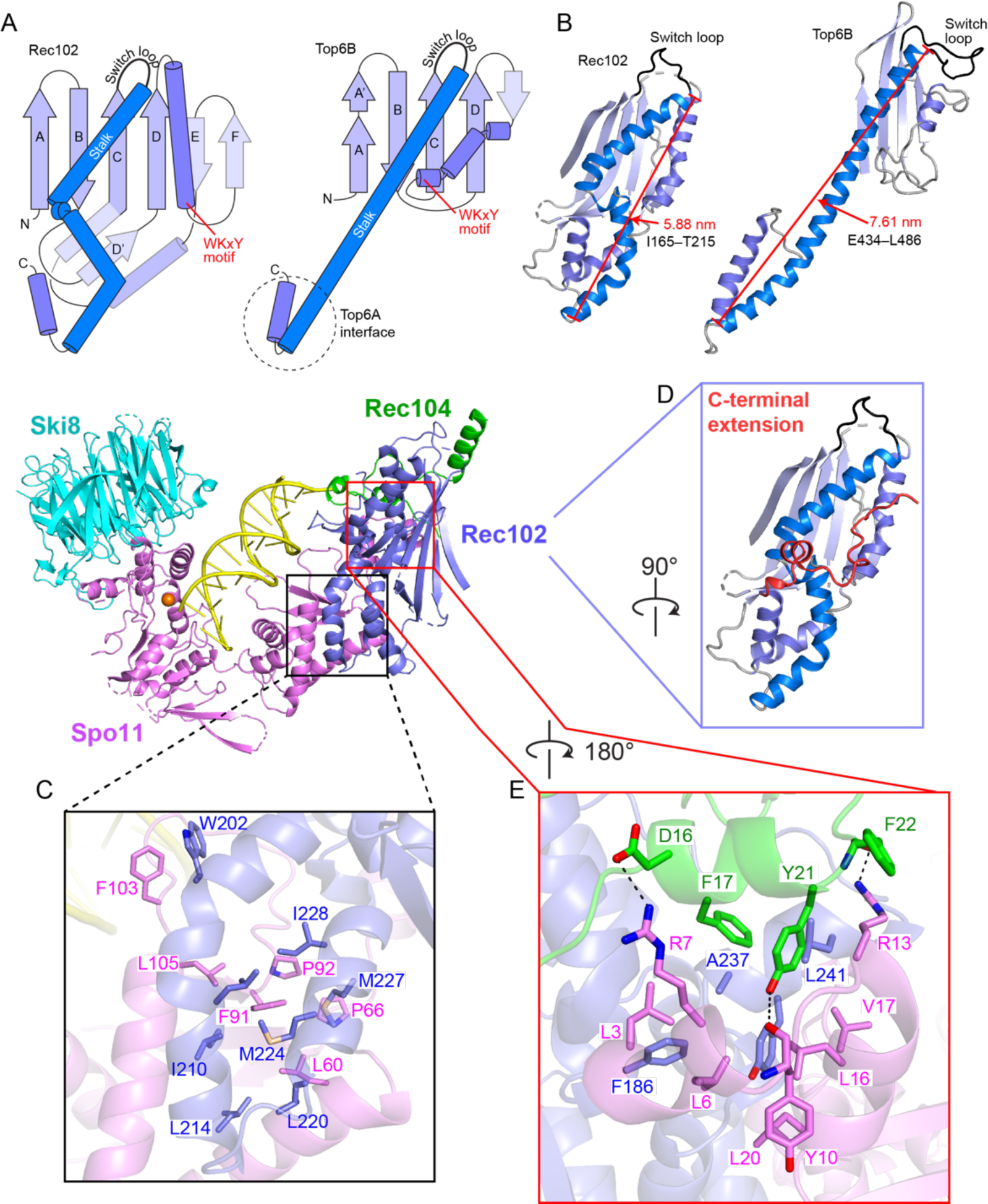
Rec102 folding and interactions with other subunits. (A) Protein topology schematics of Rec102 and Top6B highlighting conserved structural elements. (B) Ribbon representations of Rec102 (this study) and Top6B (PDB: 2zbk)^4^, with the same color settings as in panel A (C–E) Details of protein-protein contacts within the gapped-DNA-bound core complex, showing hydrophobic interactions between Rec102 and Spo11 (C); the C-terminal extension of Rec102 (red, D); and intermolecular hydrogen bonds and hydrophobic interactions between the N-terminal extension of Spo11 and Rec102 and Rec104 (E).

In Top6B, the central β sheet is a scaffold supporting the stalk helix and the GHKL and H2TH domains ^4,5,11^. Rec102 has a similar β sheet, with the first four strands corresponding to previously identified segments of homology with Top6B ^13^ (**Fig. 3A**, strands labeled A–D, and **Fig. S7A,B**). Nearly all of the residues in these strands are encoded in *REC102* exon 1, which is essential for Rec102 interaction with Rec104 and for function in vivo ^23^. The sheet is extended by one (Top6B) or two (Rec102) additional β strands, but these arise differently in the sequence: the short strand in Top6B is immediately after strand D, but the two longer strands in Rec102 instead occur after a conserved helix (**Fig. 3A**).

Another highly conserved element is a tryptophan within a motif that has the sequence WKxY in the archaeal proteins ^13^, and that starts at the end of a set of short α helices and continues into the subsequent turn (**Fig. 3A and S7B**). The lysine in this motif is part of a putative DNA binding surface in Topo VI that is critical for activity ^24^. The tryptophan is in the hydrophobic core of *Saccharolobus* Top6B, packing against the stalk and against hydrophobic residues at the beginning of strand C (**Fig. S7C**). The tryptophan is nearly invariant in eukaryotes as well, but the rest of the motif is highly variable (W_91_EEQ in Rec102) (**Fig. S7B**). In Rec102, W91 packs against the equivalent of the stalk helix, analogous to Top6B, but contacts different strands in the β sheet: I59 in strand D (β4) and L113 in strand E (β6) (**Fig. S7C)**. In the core complex structure, this motif is ≥16 Å away from the nearest DNA segment, so it does not appear to contribute to DNA binding in the end-bound complex. However, our findings do not exclude a possible role in pre-DSB binding by a core complex dimer to an intact DNA duplex.

The third conserved element, the switch loop, is a 16-residue sequence in *Saccharolobus* Top6B that connects strand C with the stalk helix (**Fig. 3A and S7B**). It contains a lysine that contacts ATP bound by the GHKL domain’s active site and that is essential for ATPase activity in Topo VI and other topoisomerases ^25–27^. Our structure confirms the earlier deduction of the position in the Rec102 primary structure equivalent to the switch loop ^13^, but the loop is smaller in Rec102 than previously appreciated (just 8 residues, S_157_KEGNYVE).

The stalk of Rec102 begins at residue I165, after the switch loop. As in Top6B, the first part of the stalk is an amphipathic helix whose hydrophobic face runs diagonally across strands D through A of the central β sheet (**Fig. 3A,B and S7D**). However, unlike the long, continuous α helix that makes up the stalk of Top6B, the equivalent region of Rec102 is distorted into four helical segments to make a kinked path that is more compact and engages in more extensive intra- and intermolecular contacts within the core complex (**Fig. 3A,B**) ^9^. We examined seven mutations along the stalk: I195A, L198A, R199A, W202A, L179A, S182A, Q183A (**Fig. S8A,B**). Of these, R199A completely eliminated both meiotic recombination in vivo (*arg4* heteroallele recombination assay) and interaction with Spo11 (yeast two-hybrid (Y2H) assay), establishing the importance of this residue, which forms hydrogen bonds with multiple residues in Spo11 (**Fig. S8C**). The rest of the mutations tested had no effect or caused at most a two-fold reduction in meiotic recombination frequency.

The Rec102 stalk winds its way to interact with the WH domain of Spo11. A helical hairpin at the end of the stalk aligns with a pair of α helices from Spo11 (**Fig. 3A,C**). The structure of this part of the Spo11-Rec102 interface resembles the computational Spo11-Rec102 model, which was earlier noted to resemble the Top6A-Top6B interface ^9^. However, Spo11 and Rec102 have a much more extensive interface than Top6A and Top6B (1,681 Å^2^ for Spo11–Rec102 vs. 958 Å^2^ for Topo VI; **Fig. S8D**), mediated by both hydrophobic and hydrogen bonding interactions (**Fig. 3C and S8C**). Mutations across the Rec102 helical hairpin (L214 and residues 220–236) were previously shown to disrupt the Y2H interaction with Spo11 and to eliminate meiotic recombination initiation in vivo ^14^. Meiotic recombination was also compromised by single alanine substitutions for interfacial residues L53, L60, and L112 in Spo11 and L207 of Rec102, but with variable effects on the Y2H interaction (**Fig. S8A,B**). Other residues in this interface were dispensable for meiotic recombination in vivo (Spo11 F103, which packs against Rec102 W202; and Spo11 L105) (**Fig. 3C and S8A,B**).

In addition to these elements that are conserved with Top6B, Rec102 sports a 34-residue C-terminal extension beginning at D231. This segment wraps around the stalk, contributes to the extended interface with Spo11, and also makes contacts with the DNA and Rec104 (**Fig. 3D** and below). Double alanine substitutions for Rec102 D231/K232 or T235/T236 were previously shown to compromise both meiotic recombination and the Spo11 Y2H interaction ^14^. Rec102 S233 and Q239 form hydrogen bonds with the Spo11 peptide backbone at G95 and L98, respectively (**Fig. S8C**). Rec102-S233A and Q239A as well as Spo11-G95A (but not L98A) severely compromise meiotic recombination in vivo and diminish the Spo11-Rec102 Y2H interaction (**Fig. S8A,B**).

There is also an extension (25 residues) of the N terminus of Spo11 before the WH domain ^16^, but the function of this extension has been unclear. This α-helical segment contacts Rec102’s C-terminal extension and stalk and the N terminus of Rec104, through hydrophobic interactions and hydrogen bonds (**Fig. 3E**). The extension on Spo11 unexpectedly places the N-termini of Spo11, Rec102, and Rec104 all in close proximity. This agrees well with the extensive crosslinking of the Spo11 N terminus with both Rec102 and Rec104, and explains why an N-terminal MBP tag on any of these proteins resulted in extra density in nearly the same position in negative-stain EM images ^14^. Several mutations in this region of Spo11 (L3A, R7D, and L20A) severely compromised meiotic recombination in vivo while only modestly diminishing Y2H interactions with Rec102 or Rec104 (**Fig. S8A,B**).

### Rec104 structure and protein interactions

Only residues 9–58 of Rec104 are visible in the structure, forming three α helices that interact with both Rec102 and Spo11 (**Fig. 1D**). Helix 1 and the preceding residues of Rec104 contact both Rec102 (strand A and the stalk) and Spo11 (N-terminal extension), primarily through backbone and side chain hydrogen-bond interactions (**Fig. 4A**). Helices 1 and 2 in Rec104, plus the intervening turn, contact the Rec102 stalk and C-terminal extension through hydrophobic interactions (**Fig. 4B**). The third α helix of Rec104 contacts the Rec102 β sheet (strands A, B and C) and switch loop through hydrophobic interactions (**Fig. 4C**). We emphasize, however, that the resolution is lower for this part of the structure (**Fig. S2D and S3D**), so the details of the interfacial contacts should be viewed cautiously.

**Fig. 4.**
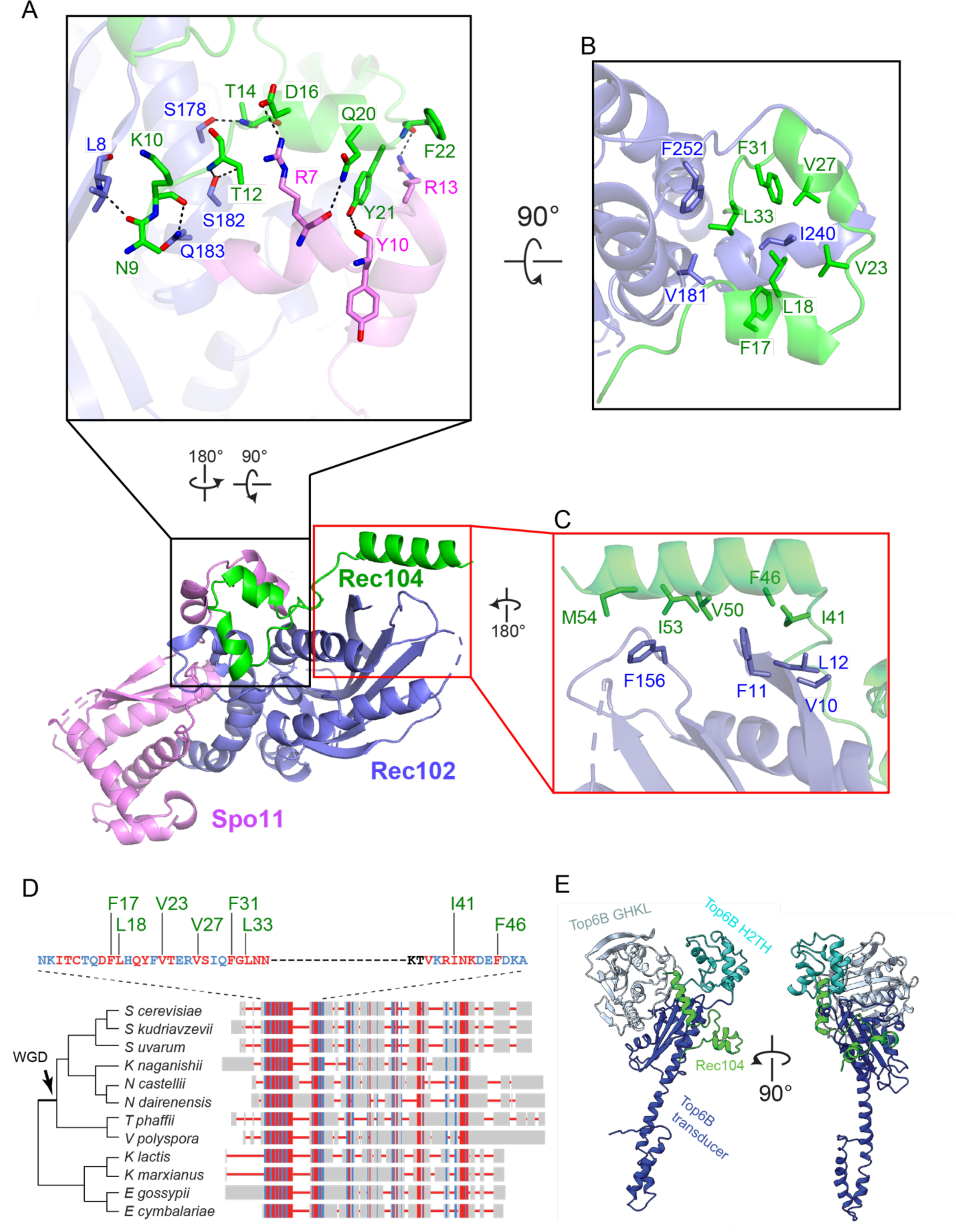
Rec104 structure and interactions with other subunits. (A–C) Details of protein-protein contacts, showing hydrogen bonds between the structured N-terminal region of Rec104 and Rec102 and Spo11 (A); hydrophobic interactions among helices 1 and 2 of Rec104, Rec102, and Spo11 (B); and hydrophobic interactions between helix 3 of Rec104 and Rec102 (C). Panels A and B show the gapped-DNA-bound core complex; panel C shows the hairpin-DNA-bound complex because the resolution was higher for this part of the structure. (D) Conservation of Rec104 in Saccharomycetaceae. A multiple sequence alignment of Rec104 sequences from twelve species was generated and visualized using COBALT^55^. Aligned residues are colored based on relative entropy, with red indicating more conserved residues. The region with strongest conservation, shown in detail above, matches the portion of Rec104 visible in the cryo-EM structures. Conserved residues mediating hydrophobic interactions are labelled in green. Topology of the cladogram is based on^58^. WGD, whole-genome duplication. (E) Position of the structured region of Rec104 relative to the domains of *S. shibatae* Top6B. The core complex was superposed with Topo VI (PDB: 2zbk)^4^ by aligning S147–F156 of Rec102 to M409–T418 of Top6B. Only Rec104 from the core complex is shown (green); the GHKL ATPase domain of Top6B is colored pale cyan, the H2TH domain is cyan, and the transducer domain is dark blue.

Previous efforts to identify Rec104 orthologs failed to find them outside of *Saccharomyces sensu stricto* species because of poor sequence conservation^28^. The structure of the core complex allowed us to revisit this because the visible parts of Rec104 correspond to a previously unrecognized conserved domain (**Fig. 4D**). We expanded the collection of Rec104 homologs by focusing on this domain, but still were only able to detect them in family Saccharomycetaceae (**Fig. 4D** and Methods). Particularly well conserved residues (**Fig. 4D**) line the interfaces with Rec102 and Spo11, including Rec104 Y21 (**Fig. 4A**); F17, L18, V23, V27, F31, and L33 (**Fig. 4B**); and I41 and F46 (**Fig. 4C**). The C-terminal portion of Rec104 that is not visible in the cryo-EM structure is poorly conserved (**Fig. 4D**).

We tested the functional importance of a number of residues across the structured region of Rec104 (**Fig. S8A,B**). The Y21A mutant was strongly defective for meiotic recombination and Y2H interaction with Spo11, but retained Y2H interaction with Rec102. The other *REC104* mutations tested had little if any effect in either assay.

Prior crosslinking experiments had suggested that Rec104 occupies a position near where the GHKL domain abuts the transducer domain in Topo VI, supporting the proposal that Rec104 replaces the GHKL domain ^1,14^. We explored this idea using our structure.

The most frequently observed crosslinks between the two proteins were between Rec104 K43 (at the beginning of helix 3) and either Rec102 K60 or K64. In the structure, the side chains of these lysines are positioned appropriately to allow crosslinks to occur, and the distances between their Cα atoms are less than the 27.4 Å maximum for the crosslinker used (19.1 Å for K43 to K64; 24.3 Å for K43 to K60) (**Fig. S8E**). The structure thus explains these crosslinking results well.

Rec102 K60 and K64 are in strand D of the central β sheet, with their side chains on the opposite face of the sheet from the stalk (**Fig. S7B and S8E**). The equivalent region in Top6B is at the interface with the GHKL domain, as previously noted ^14^. To compare the position of Rec104 to the domains of Top6B, we superimposed the core complex structure with a crystal structure of Top6B ^4^ by aligning Cα atoms of strand C from each protein. In the view of Top6B shown in **Fig. 4E** (right), the Top6B GHKL domain lies behind and to the right of the transducer domain’s central β sheet and the H2TH domain is positioned to the upper left, above the loop between strands A and B. In the superimposed ensemble, Rec104 wraps around the transducer domain, with helices 1 and 2 lying in front of the stalk and helix 3 lying against the back side of the upper corner of the β sheet (**Fig. 4E**). The structured elements of Rec104 are thus strikingly distinct from all of the domains of Top6B.

Although we do not know the disposition of the C-terminal two-thirds of Rec104 because it is not visible in the structure, the end of helix 3 points toward the location of the Top6B GHKL domain (**Fig. 4E**). Moreover, Rec104 lysines 61, 65, 78, and 79 (which lie just outside the structured segment) were found to crosslink to Rec102 K60, K64 and K79 ^14^, all three of which are in proximity to the position occupied by the GHKL domain in Top6B.

We conclude that the structured part of Rec104 is not equivalent to any of the domains in Top6B, but that the unstructured C-terminal part of Rec104 is near or in the GHKL domain position, consistent with the previous proposal about Rec104’s location ^14^. However, it remains unclear whether the unstructured portion of Rec104 evolved from the GHKL domain or is a wholly unrelated fold that replaced it.

### Protein-DNA contacts and the specificity of DNA binding

We can trace the 15-bp duplex of one hairpin element of the gapped DNA, plus overhang nucleotide A39 including its 5′-phosphate (**Fig. 2B,C**). An electrostatic surface representation of the core complex identified three positively charged patches that line the inner face of the DNA binding channel and mediate non-sequence-specific protein-DNA contacts (**Fig. 5A**).

**Fig. 5.**
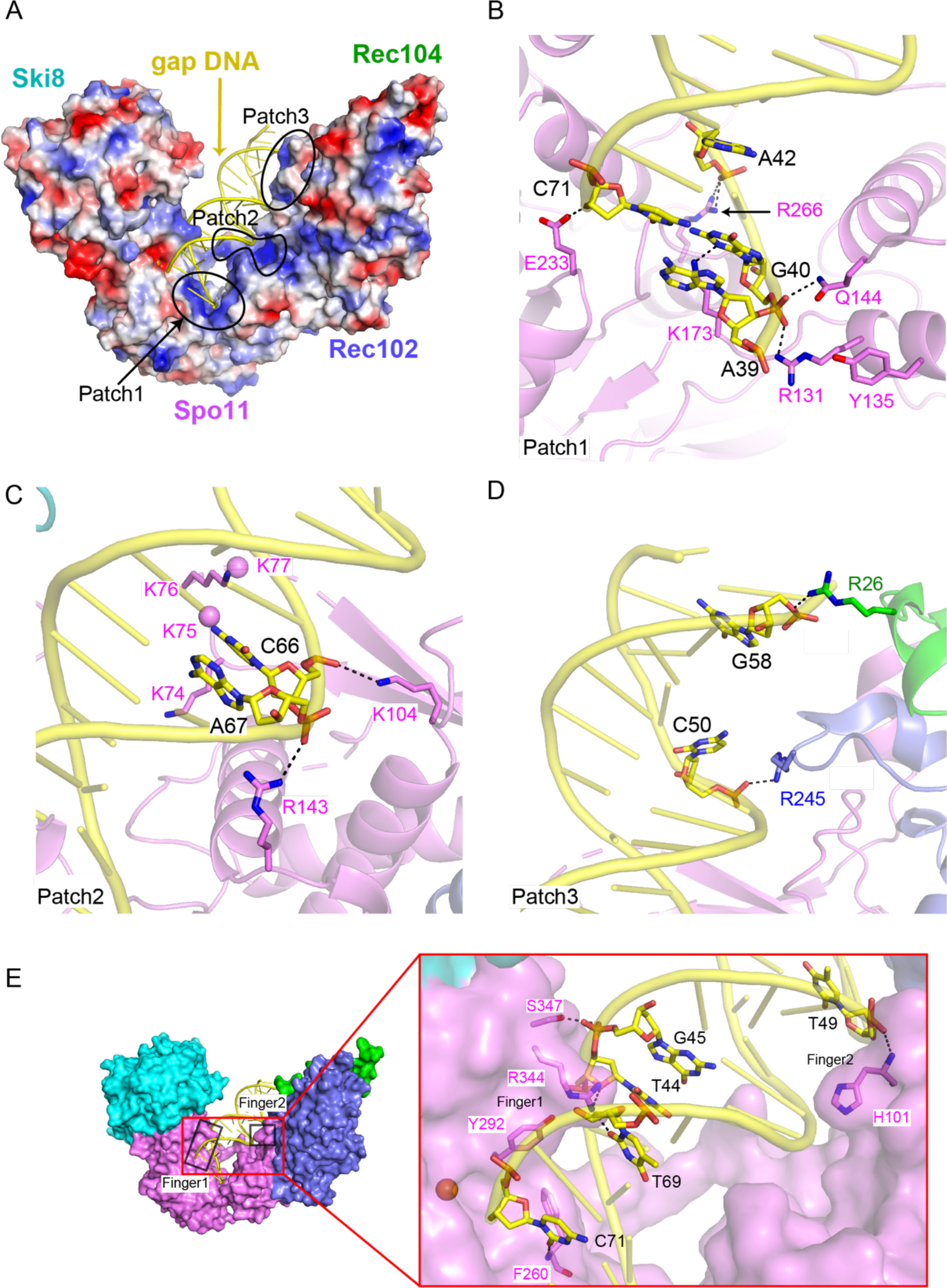
Protein-DNA contacts in the gapped-DNA-bound core complex. (A) Electrostatic surface representation of the protein subunits. (B–D) Details of protein-DNA contacts associated with patch 1 (B), patch 2 (C) and patch 3 (D). Side chain densities that could not be traced are shown as balls. (E) Details of protein-DNA contacts associated with fingers 1 and 2.

Patch 1 anchors overhang nucleotide A39 and its adjacent duplex segment in the pocket between the WH and Toprim domains of Spo11. Specifically, R131 on the second α helix of the WH domain forms a hydrogen bond with the 5′ phosphate of G40, which is the bridging phosphate between the first base of the duplex (G40) and the beginning of the 5′ overhang (A39) (**Fig. 5B**). This interaction by R131 stabilizes the orientation relative to the DNA of the catalytic tyrosine Y135, which is on the same α helix (**Fig. 5B**). In addition, the third α helix of the WH domain inserts into the major groove of the DNA, from which the side chain of residue Q144 interacts with the same phosphate as R131 (**Fig. 5B**), while R143 interacts with the phosphate backbone of the opposite strand (**Fig. 5C**). Further, residue K173 at the beginning of the flexible linker connecting the WH and Toprim domains makes one of the rare contacts of Spo11 with a base, forming a hydrogen bond with the N-3 position of G40 (**Fig. 5B**). This contact is interesting because G is strongly favored at this position for natural Spo11 cleavage events in vivo^29^. Toprim domain residues in patch 1 include R266, which forms a hydrogen bond with the phosphate of A42, and E233, which forms a hydrogen bond with the 3′-terminal hydroxyl group of C71 (**Fig. 5B**).

These interactions of patch 1 with DNA explain patterns of protein sequence conservation as well as prior experimental findings in vivo and in vitro. Specifically, R131, Q144, R266, and E233 are invariant in Spo11 and Top6A relatives, and K173 is a basic residue in nearly all eukaryotic homologs but is sometimes aspartate in Top6A (**Fig. S9A**). Moreover, the direct contacts of R131 and Q144 with the 5′ phosphate at the beginning of the duplex and of E233 with the 3′ hydroxyl explain the strong selectivity of binding to DNA with a free 3′-OH end and 5′ ssDNA overhang ^14^. R131 and E233 are essential for Spo11 function in vivo ^7^; mutating K173 to alanine reduced DNA binding in vitro and delayed and reduced DSB formation in vivo^14^; and mutating E233 to alanine reduced DNA binding in vitro by ∼10-fold ^14^.

Patch 2 is in the central segment of the DNA-binding channel and comprises residues from the Spo11 WH domain (**Fig. 2A and 5A**). The basic K_74_KKK loop surrounds the T63-T64-A65 containing strand, while K104 forms hydrogen bonds with the phosphate of C66 (**Fig. 2A and 5C**). The density is weak for the K_74_KKK loop, with K74 and K76 directed towards the DNA sugar-phosphate backbone. This loop is part of a beta hairpin (the “wing” in the WH domain) and is a prominent site of cleavage by hydroxyl radicals produced by iron chelating moieties placed at nearby positions on the DNA ^14^, but is highly variable in length and sequence in Spo11 family members (**Fig. S9A**).

Patch 3 is on the outer segment of the DNA-binding channel (**Fig. 2A and 5A**). It includes R245 from the C-terminal extension of Rec102, which forms a hydrogen bond with the phosphate of C50, while R26 of Rec104 forms hydrogen bonds with the phosphate of G58 (**Fig. 5D**).

In addition to these positively charged DNA-binding patches, we also identified two sets of residues (“fingers”) projecting from Spo11 into the minor groove, stabilizing the bound hairpin DNA segment in the complex. Finger 1—consisting of F260, Y292, and R344 from the Toprim domain—tracks along the minor groove of the first five base pairs of the duplex, with R344 forming base-specific hydrogen bonds with the O-4’ of T44 and O-2 of T69, further stabilized by hydrogen bonding of S347 with the phosphate of G45 (**Fig. 5E**). The F260 main chain nitrogen also forms a hydrogen bond with the O-2 of base C71. F260 is a bulky hydrophobic residue in Spo11 orthologs but glutamine in most Top6A proteins (**Fig. S9A**). The minor groove binding by finger 1 is particularly striking because we previously noted that the base composition bias around Spo11 cleavage sites in vivo is consistent with a tendency toward a relatively wide and shallow minor groove across precisely this region^29^. An F260A mutation dramatically changes DSB site preference in vivo and reduces DNA-binding affinity in vitro ^14^, while mutating Y292 to arginine disrupts DSB formation in vivo ^7^. Finger 2 consists of H101 from the WH domain, which forms a main-chain amide hydrogen bond with the phosphate of T49 and inserts its imidazole ring into the DNA minor groove (**Fig. 5E**).

### Metal ion binding

Type IIA topoisomerases (eukaryotic Topo II and bacterial gyrase and Topo IV) are thought to use a two-metal mechanism involving divalent cations bound by acidic residues of the Toprim domain ^30–32^. In this model, metal ion A interacts with both bridging and non-bridging oxygens of the scissile phosphate and plays a direct role in transesterase catalysis, while metal ion B interacts with an adjacent (non-scissile) phosphate and plays a structural role in stabilizing protein-DNA interactions (**Fig. 6A**). Comparatively little is known about the detailed roles of metals in type IIB enzymes, aside from their requirement for DNA cleavage by Topo VI ^3,33^.

**Figure 6.**
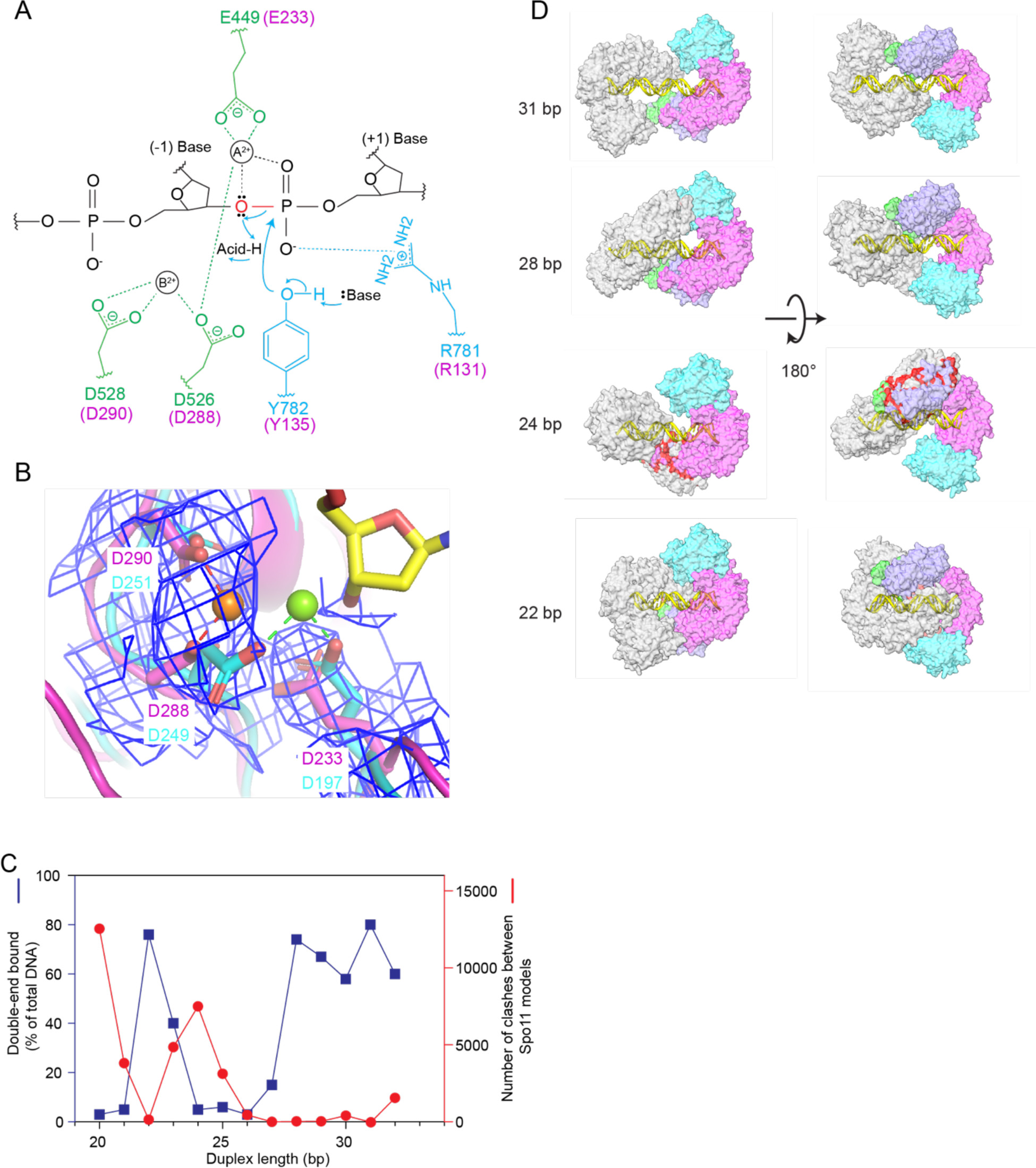
Metal ion binding and models of higher order protein-DNA complexes. (A) Proposed two-metal-ion mechanism for catalysis of strand cleavage by type IIA topoisomerases (schematic adapted from Schmidt et al.^30^). Amino acids in the active site of *S. cerevisiae* Topo II are indicated, with metal-binding residues in the Toprim domain colored in green and residues from WH domain colored in blue. Corresponding Spo11 residues are colored in magenta. The general base and acid are unknown. (B) Superposition of the metal-binding pockets of Spo11 (magenta) and *Methoanocaldococcus jannaschii* Top6A (cyan, PDB: 1d3y^6^). Only the metal ion and the side chains of the indicated residues from Top6A are shown. The magnesium in the Spo11 cryo-EM structure is coordinated by D288 and D290, while the magnesium in Top6A is coordinated by D249 and E197, which are equivalent to D288 and E233 in Spo11. The electron density map around the Spo11 triads and Mg^2+^ is shown in blue mesh; note the lack of detectable density at the position of the metal (green ball) bound by Top6A (C,D) Models for two Spo11 core complexes bound to opposite ends of DNA duplexes of varying lengths. The plot in (C) compares the number of clashes between the two core complexes as a function of duplex DNA length (red) with direct measurement of the ability to form double-end-bound complexes in EMSA experiments (blue; data from ^35^). The blue points show the fraction of protein-DNA complexes that have both DNA ends bound. Panel D shows example models of double-end bound core complexes with DNA duplexes of 22 bp, 24 bp, 28 bp and 31 bp (lengths do not include the ssDNA overhangs at each end). One core complex is colored as in Fig.1D. The other core complex is colored in gray. Clashes are colored red.

We observed density in both cryo-EM structures consistent with a single metal ion bound by D288 and D290 of Spo11 (**Fig. 6B**). This is most likely Mg^2+^, which was present when the protein-DNA complexes were assembled. When the Spo11 metal-binding pocket was compared with the crystal structure of *Methanocaldococcus jannaschii* Top6A ^6^, the trio of highly conserved acidic residues was highly congruent but the single Mg^2+^ bound by each protein was in a different position, bound by E197 and D249 in Top6A (equivalent to Spo11 E233 and D288) (**Fig. 6B**). By comparison to analogous residues in a structure of yeast Top2 ^30^, we infer that site A is the one occupied in the Top6A structure, whereas site B is the one occupied in our DNA end-bound Spo11 structure (**Fig. 6A,B**). In type IIA enzymes, site A has a higher affinity for metal than site B, but is thought to rely on the scissile phosphate for two of the metal coordination contacts ^32^. If the same is true for Spo11, the absence of an equivalent of the scissile phosphate at the 3′ end in our structure may explain why Mg^2+^ is not stably bound at site A. The observed binding in site B is consistent with the presumed structural role for the metal in this position. Occupancy of site B is consistently observed in post-cleavage structures of eukaryotic and bacterial topoisomerases ^30,34^, supporting the conclusion that our structure of the non-covalently bound Spo11 core complex mimics the post-cleavage state.

These findings provide a framework for understanding the sequence conservation and functional importance of the metal-binding acidic residues in Spo11. E233 and D288 are invariant in Spo11 and Top6A proteins (**Fig. S9A**) and are essential for DSB formation in vivo in yeast ^7^. Extrapolating from type IIA enzymes, we propose that the essentiality of both residues traces to their directly coordinating the catalysis-critical Mg^2+^ in site A (**Fig. 6A,B**). By contrast, D290 is mostly but not strictly conserved (**Fig. S9A**), and it is dispensable for DSB formation in vivo: substitution with asparagine has little effect on recombination activity in *S. cerevisiae* while alanine substitution allows DSB formation but causes a cold-sensitive phenotype for untagged Spo11 and a ∼5-fold decrease in DSBs when combined with a particular C-terminal epitope tag^7^. It may be that the partial contribution of this residue to Spo11 function reflects that it contributes to metal binding only in the non-catalytic site B (**Fig. 6A,B**).

### Modeling higher order complexes

#### Spacing between adjacent DSBs

The cryo-EM structure sheds light on previously observed spatial patterns of Spo11-DNA interactions both in vivo and in vitro. Spo11 usually cuts a chromosome only once, but multiple Spo11 complexes can introduce two or more DSBs close together on the same DNA molecule ^35–37^. When such double-cutting occurs, it has a preferred spacing between DSBs with a minimum of ∼33 bp (measured from the center of each DSB’s 5’ overhang, corresponding to 31 bp of duplex DNA plus the overhangs) and increasing in steps of ∼10 bp. The 10-bp periodicity has been proposed to reflect a geometric constraint in which adjacent Spo11 dimers are co-oriented with their active sites facing in the same direction ^35,37^. In this model, the observed minimum distance reflects steric constraints that limit how close adjacent co-oriented Spo11 complexes can be to one another. EMSA experiments with yeast Spo11 core complexes support this interpretation: both 5′-overhang ends of a DNA substrate are readily bound if the duplex DNA segment between the overhangs is ≥28 bp, but duplex lengths of 24–27 bp can only be bound at a single end, while reducing the duplex length even further (22–23 bp) allows core complexes to again bind at both ends ^35^ (**Fig. 6C**). These results suggest that steric clashes that preclude double-end binding at 24–27 bp are relieved if the two DNA ends are rotated relative to one another. (Note that double-end binding by monomeric core complexes in vitro likely mimics how close two adjacent Spo11 dimers can be on an intact DNA molecule before cleavage.)

These striking patterns of DNA cleavage in vivo and DNA binding in vitro are well explained by the extensive left-handed wrap of the core complex around the DNA in the cryo-EM structure. Two Spo11 core complexes could be modeled on each end of a 28-bp DNA duplex with essentially no steric clashes, but modeling on a shorter DNA segment (24 bp) resulted in substantial overlap between the Rec102 and Rec104 moieties of the two core complexes (**Fig. 6C,D**). Consistent with EMSA data, the clashes could be almost entirely resolved by shortening the DNA still further to 22 bp, which rotates the core complexes relative to one another and allows them to interdigitate (**Fig. 6C,D**). Moreover, a 31-bp duplex is the shortest distance that can accommodate a pair of core complexes that is both closely co-oriented and lacking in steric clashes (**Fig. 6D**). This agrees well with in vivo spacing because a 31-bp duplex corresponds to a center-to-center DSB distance of 33 bp, i.e., the minimum preferred distance for double cuts ^35,37^.

#### Structure-based model of a pre-DSB Spo11 dimer

We further attempted to model a catalytically competent Spo11 dimer by docking two copies of the monomeric DNA-bound core complex together. If the two DNA copies are aligned to overlap the two nucleotides of the 5’ overhangs to make a roughly B-form duplex, the WH domain from each Spo11 clashes sterically with the Toprim domain from the other (**Fig. 7A**). In addition, the non-overhang strand from one complex clashes with the C-terminal part of α helix 8 of Spo11 from the other complex, and the overhang strand collides with a loop near finger 1 (**Fig. 7B**). Thus, the configurations of both protein and DNA in the cryo-EM structure are incompatible with a plausible structure of a pre-DSB complex on B-form DNA.

**Figure 7.**
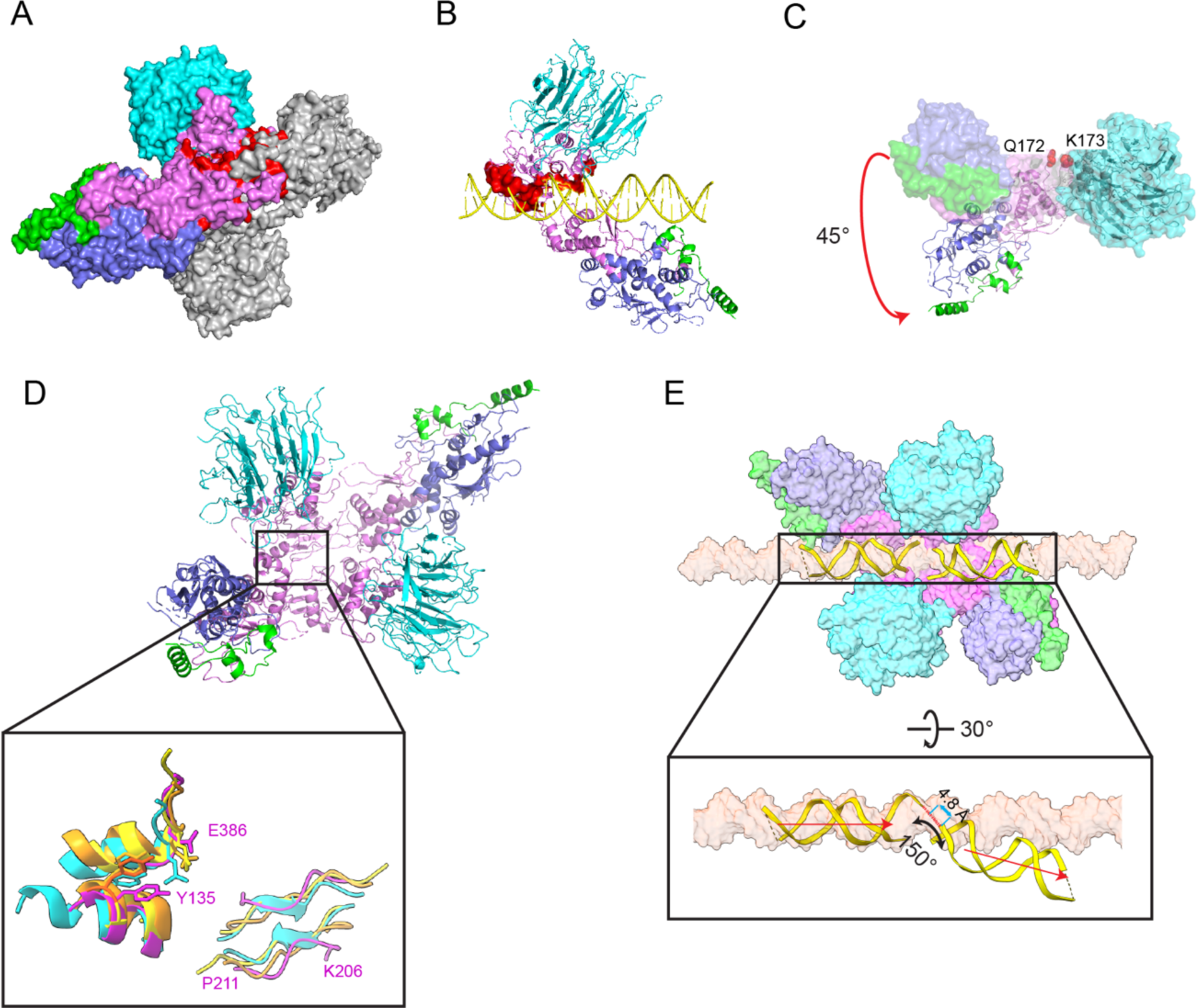
Model of a pre-DSB Spo11 dimer. (A) Predicted Spo11–Spo11 clashes when core complexes are docked together on a B-form DNA. Two copies of the DNA-bound core complexes were aligned onto a B-form DNA via the first two nucleotides of their 5’ overhangs. One Spo11 core complex is colored as Fig. 1D, and the other is colored in gray. The protein clashes are colored in red. DNA is not shown. (B) Predicted clashes between protein and DNA strands. The clashing regions are highlighted in red surface representation. (C) Rigid body motion of the Spo11 WH domain relative to the Toprim domain. The Spo11 core complex cryo-EM structure is shown in a surface representation. The ribbon diagram shows the hypothetical configuration of the core complex if the WH and Toprim domains of Spo11 are superposed separately (separated between Q172 and K173, see Methods) onto their cognate domains in *S. shibatae* Top6A from the crystal structure of the Topo VI holoenzyme^4^. (D) Comparison of parts of the Spo11 dimer interface (purple) with the equivalent Top6A dimer interfaces from different species (cyan, *M. jannaschii*; orange, *Methanosarcina mazei;* yellow, *S. shibatae*.). (E) Deformation of DNA in the hypothetical pre-DSB dimer model. Proteins of two core complexes are shown in a surface representation and the two DNA duplexes are shown in ribbon diagram with their positions relative to the Spo11 WH domains preserved from the cryo-EM structures. A surface representation of a B-form DNA is included for comparison.

Because Spo11 core complexes are flexible in solution^14^, we asked if simple rigid body motions of their domains could resolve the clashes. To answer this, we separately aligned the Spo11 WH domain (together with the DNA, Rec102, and Rec104) and the Toprim domain (together with Ski8) to the cognate domains of Top6A in the *Saccharolobus shibatae* Topo VI dimer structure ^4^ **(Fig. S9B**), because the polypeptide segment between the domains is thought to be a flexible linker^6,14^. This alignment results in rotation of the WH domain 45° relative to the Toprim domain and, notably, eliminates all of the steric clashes (**Fig. 7C**). The model also matches segments of Spo11 well with the two major Top6A–Top6A interfaces in Topo VI dimer structures ^4–6^: a pseudo-continuous β-sheet on the underside of the Toprim domain away from the DNA (K206 to P211 from one Spo11 interacting with its match on the other Spo11), and an interface between the WH domain of one Spo11 near the catalytic tyrosine and a C-terminal region of the other Spo11 that includes an invariant residue E386 that is brought in close proximity to the catalytic tyrosine (**Fig. 7D and S9A**). This model thus appears to provide a plausible representation of a dimeric pre-DSB protein-DNA complex.

The model predicts an intriguing deformation of the DNA. We preserved the relationship of the DNA to the WH domain because this also preserves the large majority of the protein-DNA contacts from the cryo-EM structure. In doing so, the overhang ends of the two DNA segments come into close proximity (∼4.8 Å separating the inferred positions of 5′ and 3′ ends of each strand), but make a V shape with a 150° angle and with the helical axes slightly offset by ∼4 Å (**Fig. 7E**). Empirical evidence for DNA deformation by Spo11 comes from the 130° angle of the DNA in our gapped double-hairpin structure (**Fig. 2D**) and by previous DNA-binding experiments that revealed bent DNA by atomic force microscopy and apparent preferential binding to bent duplex DNA (inferred from higher affinity for binding 100-bp vs. 400-bp minicircles)^14^. Topo VI and other type II topoisomerases bend DNA before cleavage^6,14,38^, so it is likely that Spo11 core complexes may do so as well. Our model thus provides a framework for understanding the structural determinants of DNA bending, for which no high-resolution structural information is currently available for any dimeric Topo VI or Spo11 complex.

## Conclusions

Ever since the discovery of the DSB-forming activity of Spo11 more than a quarter century ago ^39,40^, there has been a notable lack of empirical structures of Spo11 and its accessory proteins. Our cryo-EM structures now provide insight into the architectures of the individual subunits and of their interfaces with each other and with the DNA. We confirm the structure and position of Ski8, uncover unexpected differences between Rec102 and archaeal Top6B proteins, and partially resolve outstanding questions about the structure and spatial disposition of Rec104. Our findings further explain the high affinity and exquisite selectivity of core complex binding to DNA ends that have a recessed 3’-OH. Finally, the structures provide a molecular framework to understand striking nonrandom patterns of DSB formation in vivo, including the biased base composition around Spo11 cleavage sites and the spacing of double cuts.

Tight binding of Spo11 to DSB ends has been proposed to cap DNA breaks and contribute to control of downstream steps in recombination ^14,20^. Although we cannot exclude the possibility that the protein-DNA complexes resolved here are rearranged significantly from the conformation after strand cleavage, we consider it likely that the end-bound structures of the monomeric core complexes mimic the products of the DSB-forming reaction. If so, our findings provide insight into the structural determinants of tight end binding and may guide a targeted search for mutants selectively defective for DSB capping.

### Limitations of the study

An important unanswered question is why the yeast core complexes are not competent to cleave DNA in vitro despite having the same minimal catalytic components found in cleavage-competent Topo VI. One issue is that the high affinity for DNA ends may preclude assembly of dimers of core complexes. Our prior work indicating a preference for binding to bent DNA duplexes ^14^ motivated our experiments with the gapped double-hairpin substrates, on the premise that the ssDNA region would provide the flexibility needed to assemble dimeric complexes that resemble the pre-DSB state. However, the majority of particles again had just a single copy of the Spo11 core complex. It remains to be determined whether inclusion of other accessory proteins such as Rec114 and Mei4 is needed to allow dimer assembly and DNA cleavage in vitro ^41–44^.

## Materials and methods

### Purification of *S. cerevisiae* Spo11 core complexes

The expression plasmids and baculoviruses were prepared as described previously^14^. The Spo11 core complex was expressed by coinfecting *Spodoptera frugiperda* Sf9 cells with a combination of viruses at a multiplicity of infection of 2.5 each as described previously^14^. Typically, 500 ml of suspension culture was collected 62 hr after infection. The cells were lysed by sonication in 25 mM HEPES-NaOH pH 7.4, 500 mM NaCl, 0.1 mM dithiothreitol (DTT), 20 mM imidazole, 1× Complete protease inhibitor tablet (Roche) and 0.1 mM phenylmethylsulfonyl fluoride (PMSF) and centrifuged at 43,000 *g* for 30 min. Cleared extract was loaded onto 1 ml NiNTA resin (Qiagen) preequilibrated with Nickel buffer (25 mM HEPES-NaOH pH 7.4, 500 mM NaCl, 10% glycerol, 0.1 mM DTT, 20 mM imidazole, 0.1 mM PMSF) then washed extensively with Nickel buffer. Core complexes were eluted in Nickel buffer containing 250 mM imidazole, then further purified on Anti-Flag M2 affinity resin (Sigma). To do so, fractions from the NiNTA elution containing protein were pooled and diluted in 3 vol. of Flag buffer (25 mM HEPES-NaOH pH 7.4, 500 mM NaCl, 10% glycerol, 1 mM EDTA) before binding to the Anti-Flag resin. Core complexes were eluted from anti-Flag resin with Flag buffer containing 250 μg/ml 3xFlag peptide (Sigma). Fractions containing protein were pooled and loaded on a Superdex 200 Increase 10/300 GL column (Cytiva) preequilibrated in 25 mM HEPES-NaOH pH 7.4, 300 mM NaCl, 5 mM EDTA, 2 mM DTT. Fractions containing protein were concentrated in 50-kDa-cutoff Amicon centrifugal filters (Millipore). Aliquots were frozen in liquid nitrogen and stored at −80 °C.

### Cryo-EM sample preparation, data acquisition, processing, model building and refinement

Purified core complexes were diluted to a final concentration of ∼0.2 mg/ml and incubated with the hairpin (5’-TAGCAATGTAATCGTCTATGACGTTAACGTCATAGACGATTACATTGC) or gapped DNA (5’-TAGGCCGTCGGCTACTAAAAGTAGCCGACGGCCGGATTAGCAATGTAATCGTCTTAAGACGATTACATTGC) substrates at a molar ratio of 1:2 in binding buffer (25 mM HEPES-NaOH, pH 7.5, 150 mM NaCl, 5 mM MgCl_2_, 2 mM DTT and 2% glycerol) for 1 hr on ice. An aliquot of the incubated mixture (3.5 μl) was applied onto glow-discharged UltrAuFoil 300 mesh R 1.2/1.3 grids (Quantifoil) at ∼4 °C. Grids were blotted for 2 s at 100% humidity at 4 °C and flash frozen in liquid ethane using a Vitrobot Mark IV (Thermo Fisher Scientific).

Images were collected on a Titan Krios G2 (FEI) transmission electron microscope operating at 300 kV with a K3 direct detector (Gatan) using a 1.064 Å pixel size at the Memorial Sloan Kettering Cancer Center. The defocus range was set from −1.0 to −2.5 μm. Movies were recorded in super-resolution mode at an electron dose rate of 20 e-/pixel/s with a total exposure time of 3 s and intermediate frames were recorded every 0.075 s for an accumulated electron dose of 53.00 e−/Å^2^.

The protocol for cryo-EM reconstruction is presented in **Fig. S2A-D** and **Fig. S3A-D**. The super-resolution movies (0.532 Å/pixel) were motion corrected and 2 × Fourier-cropped using MotionCor2^45^. Contrast transfer function parameters were estimated by CTFFIND-4^46^. All other steps of image processing were performed by Relion 3.0^47^ and Cryosparc v3.3.0^48^.

For hairpin-bound core complexes, after blob picking from 2,338 images without reference, a total of 1,829,931 particles extracted after 3-pixel binning were applied to 2 rounds of 2D classification. All the particles were applied to multiple rounds of heterogeneous refinement by using the 4.8 Å model generated from Relion 3.0. The final 128,787 particles were polished and yielded a reconstruction electron microscopy map with an average resolution of 3.66 Å. All reported map resolutions are from gold-standard refinement procedures with the Fourier shell correlation cutoff being 0.143 criterion (**Fig. S2**).

For gapped-DNA-bound core complexes, after blob picking from 5,521 images without reference, a total of 7,804,373 particles extracted after 4-pixel binning were applied to the first round of 2D classification. 6,930,069 particles selected from 2D classification were applied to multiple rounds of 3D classification. One of the 3D classes with good secondary structural features and the corresponding 1,314,980 particles were reextracted without binning. The selected particles were imported to new initial models and hetero refinement, and 548,674 particles were selected from the best class for homogeneous refinement. After non-uniform refinement and local refinement, we obtained an electron microscopy map with an average resolution of 3.29 Å. All reported map resolutions are from gold-standard refinement procedures with the Fourier shell correlation cutoff being 0.143 criterion (**Fig. S3**).

Model building was performed manually based on the cryo-EM density map and a computed model by AlphaFold2^10^ using Coot^49^. Atomic coordinates were refined against the map by real-space refinement in Phenix^50^ by applying geometric and secondary structure restraints. Details of data collection, image processing and model building are listed in **Table S1** and shown in **Fig. S2 and S3**. All figures were prepared by PyMol (https://pymol.org/2/), UCSF Chimera^51^, or UCSF ChimeraX^52^.

### DNA substrates and EMSA assays

EMSAs used the same DNA substrates as in the cryo-EM structures. Both the hairpin and gapped substrates were assembled by self-annealing in annealing buffer (100 mM NaCl, 10 mM Tris-HCl pH 8, 1 mM EDTA). Substrates were 5’ end-labeled with [*γ*-^32^P]-ATP (Perkin Elmer) and T4 polynucleotide kinase (NEB) and purified by native polyacrylamide gel electrophoresis. Binding reactions (10 µl) were carried out in 25 mM Tris-HCl pH 7.5, 7.5% glycerol, 100 mM NaCl, 2 mM DTT, 5 mM MgCl_2_ and 1 mg/ml BSA with 0.1 nM DNA. Complexes were assembled for 30 min at 30 °C and separated on 6% polyacrylamide DNA retardation gels (Invitrogen). Gels were dried and analyzed by phosphorimaging (Fuji).

### Modeling higher order assemblies of Spo11 core complexes

All modeling was done in ChimeraX^53^. The Spo11 double-end binding complex was modeled by aligning the DNA from a Spo11 core complex structure to each end of a 34-bp B-form DNA. To generate models with different lengths of DNA, the core complex at one end was fixed, while the other Spo11 core complex was moved along the B-DNA path by the specified number of base pairs.

To model a potential pre-DSB assembly containing two core complexes bound to B-form DNA, the individual DNAs in two copies of the cryo-EM structure were each aligned to a B-form DNA segment such that the first nucleotide in the 5′ overhang from one DNA was 1 nucleotide away from the 3′ end of the other DNA.

To model a pre-DSB dimer based on a Topo VI dimer, the cryo-EM structure was separated into two parts: one included residues 1–172 of Spo11 plus Rec102 and Rec104, and the other included the remaining C-terminal part of Spo11 plus Ski8. The two parts were aligned separately to the cognate domains of the *S. shibatae* Top6A subunit from a crystal structure of the Topo VI holoenzyme (PDB: 2zbk)^4^. The alignment of N-terminal part of Spo11 resulted in an rmsd of ∼1.2 Å over 32 atom pairs; alignment of the C-terminal part of Spo11 resulted in an rmsd of ∼1.1 Å over 70 atom pairs. The session files are available at doi: 10.17632/dx3hx827fp.1.

### Yeast strains and targeting vectors

All yeast strains are from the SK1 background (**Table S2**). All the vectors are generated based on pSK275, pSK276, pSK282, pSK293, pSK305, pSK310 for both yeast two-hybrid assays and heteroallele recombination assays (**Table S3**).

### Yeast two-hybrid assays

Y2H vectors were transformed separately into haploid strains SKY661 and SKY662 and selected on appropriate synthetic dropout medium. Strains were mated and streaked for single diploid colonies on medium lacking tryptophan and leucine. Single colonies were grown overnight in selective medium containing 2% glucose. Cells were lysed and quantitative β-galactosidase assay was performed using Yeast β-Galactosidase Assay Kit following the manufacturer’s protocols (Thermo Scientific). For Y2H experiments in meiotic conditions, after overnight culture in selective medium with glucose, cultures were washed twice then incubated with 2% KOAc for 20 h to induce meiosis.

### Heteroallele recombination assays

Heteroallele recombination assays were done in diploid strains heterozygous for the *arg4-Bgl* and *arg4-Nsp* alleles^54^ along with homozygous deletion mutations for *spo11*, *rec102* or *rec104*. Complementation test plasmids expressing wild type or mutant versions of Spo11, Rec102, or Rec104 (**Table S3**) were introduced by lithium acetate transformation. Three independent colonies were cultured overnight on synthetic complete medium lacking tryptophan and leucine to select for the test plasmid, then transferred to YPA (1% yeast extract, 2% peptone, 1% potassium acetate) for 13.5 h before meiotic induction in 2% potassium acetate. After at least 3 h in meiosis, appropriate dilutions of cultures were plated on synthetic complete medium lacking arginine to measure the frequency of Arg+ recombinants and on YPD (1% yeast extract, 2% peptone, 2% dextrose) to measure colony forming units.

### Phylogenetic analyses

Protein-coding sequences of Rec104 from different fungal species were identified using NCBI BLAST. Conserved regions were identified using a subset of twelve species and visualized using COBALT^55^. For the analysis of structural conservation in Rec102 and Top6BL homologs, Alphafold2 structure predictions were downloaded from the Alphafold2 database^56^. A BLAST search for a Rec102 homolog in *Sordaria macrospora* identified a predicted structure (AF-F7VPQ4-F1, NCBI XP_003350816.1) which lacks one of the conserved beta strands and the characteristic WKxY motif. Inspection of the genomic locus suggests that a splice site is misannotated; correcting this results in a longer second exon, adding the missing sequence. We used the new sequence to model the structure using Colabfold^57^.

### Data Availability

The atomic coordinates and cryo-EM density maps for the hairpin DNA bound to Spo11 core complex (PDB: 8URU; EMD: EMD-42501) and for gapped DNA bound to Spo11 core complex (PDB: 8URQ; EMD: EMD-42497) have been deposited in the Research Collaboratory for Structural Bioinformatics Protein Data Bank and Electron Microscopy Data Bank, respectively.

## Acknowledgments

This article is subject to the Open Access to Publications policy of the Howard Hughes Medical Institute (HHMI). HHMI lab heads have previously granted a nonexclusive CC BY 4.0 license to the public and a sublicensable license to HHMI in their research articles. Pursuant to those licenses, the author-accepted manuscript of this article can be made freely available under a CC BY 4.0 license immediately upon publication.

We thank members of the Keeney and Patel laboratories for discussions and experimental advice, and Jason De La Cruz (Memorial Sloan Kettering (MSK)) for support for cryo-EM data collection. MSK core facilities are supported by National Cancer Institute Cancer Center support grant P30 CA08748. KL is a Damon Runyon Fellow supported in part by the Damon Runyon Cancer Research Foundation (DRG-[2389-20]). MA was supported in part by an EMBO Long Term Fellowship (ALTF 905-2019). This work was supported by NIH grant R01 HD110120 (to SK and DJP); an MSK Basic Research Innovation Award (BRIA, to SK and DJP); a postdoctoral BRIA (to MA); Leukemia and Lymphoma SCOR 7021020 grant and Maloris Foundation (to DJP); and NIH grant R35 GM118092 (to SK).

## Author contributions

CCB, SK, and DJP conceived the project; YY, JW, KL, ZZ, MA, CCB, SK and DJP designed experiments; YY, JW, KL, ZZ, MA, CCB and SP performed experiments; YY, JW, KL, ZZ, MA, CCB, SK and DJP analyzed data; SK and DJP supervised the research; KL, MA, SK and DJP secured funding; YY, JW, KL, SK and DJP wrote the paper with input from ZZ and MA; all authors edited the manuscript.

