## Supplemental Figures and Tables for "Cryo-EM structure of the Spo11 core complex bound to DNA"

**This pdf contains:**

Supplemental Figures S1-S9

Supplemental Tables S1-S3

### SUPPLEMENTAL FIGURES

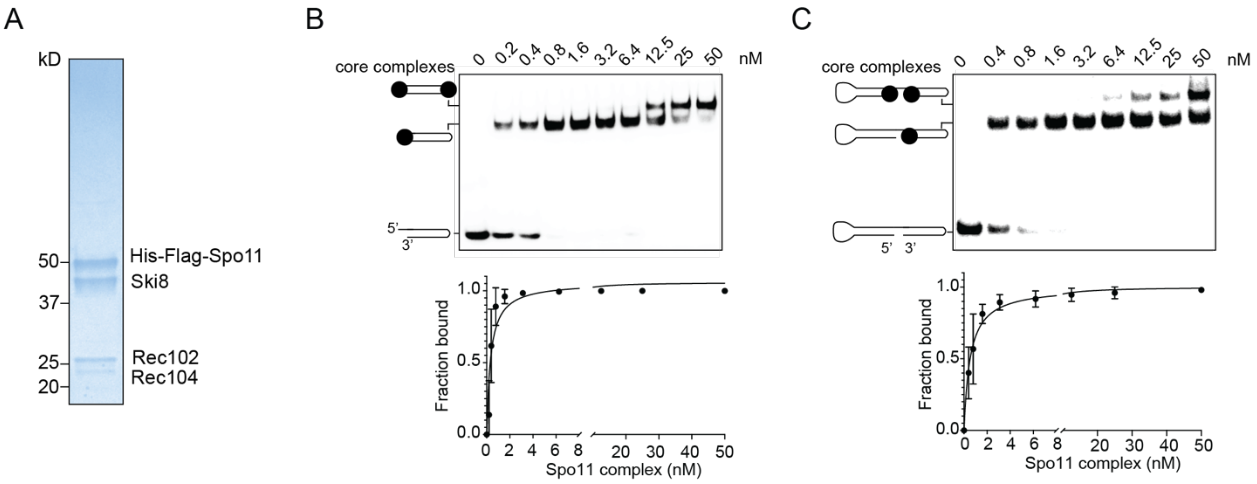


**Fig. S1. Spo11 core complex purification and EMSA assays.**
(A) SDS-PAGE gel of purified core complex (~1 µg), stained with Coomassie.
(B,C) EMSA assays and quantification of core complex binding to the hairpin DNA substrate (B) and the gapped DNA substrate (C). In panel C, the position of the second core complex that gives the slowest migrating band is unknown; it could be at the nick location as shown or it could associate with one of the hairpin ends. Error bars indicate mean ± SD of three replicates.

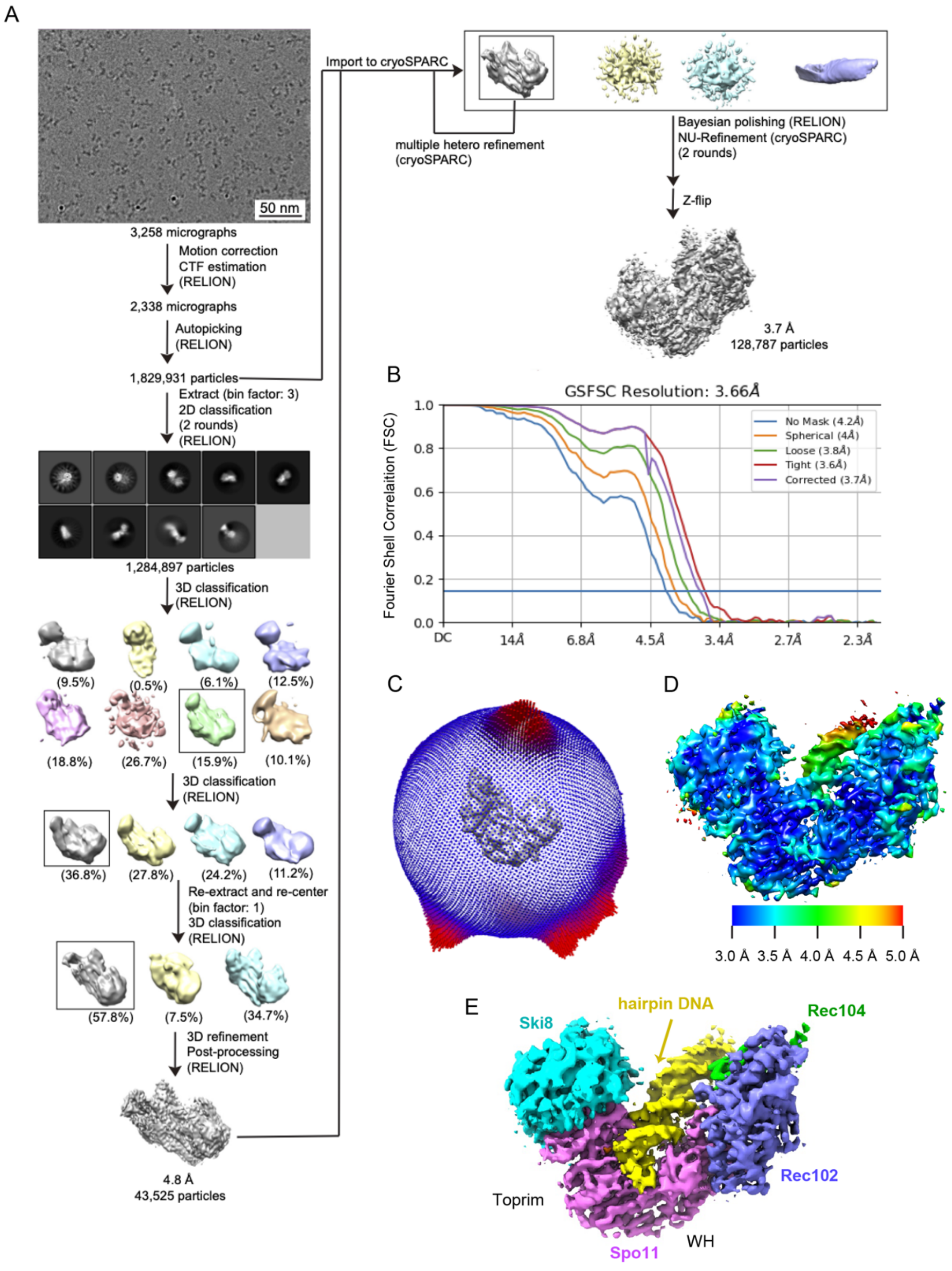


**Fig. S2. Cryo-EM reconstruction of the core complex bound to hairpin DNA.**
(A) Flow chart of cryo-EM image processing.
(B) Global Fourier shell correlation (FSC) curves. The overall cryo-EM map resolution is 3.7 Å with FSC set at 0.143.
(C, D) Euler angle distribution (panel C) and final 3D reconstructed map colored according to local resolution (panel D).
(E) Density map.

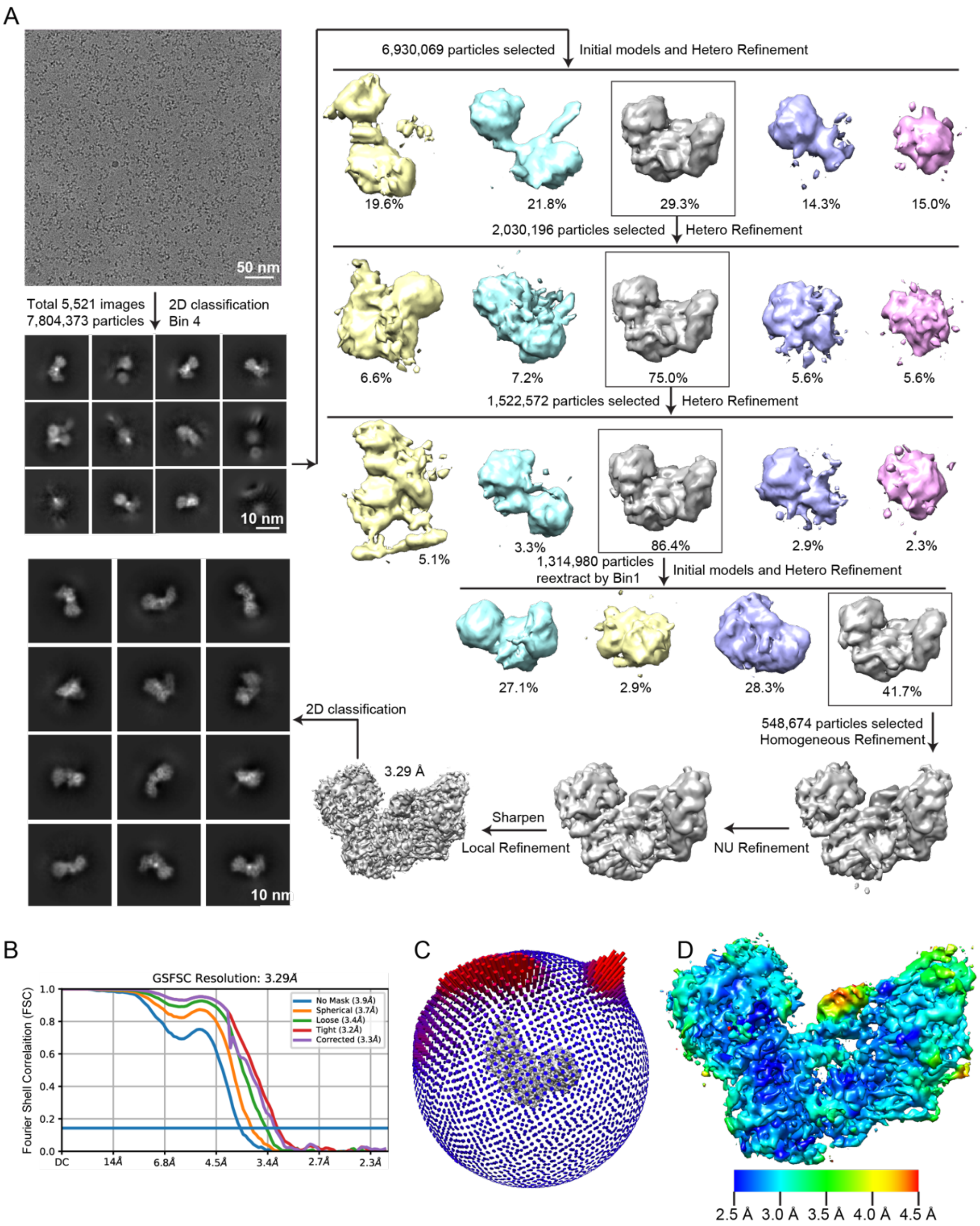


**Fig. S3. Cryo-EM reconstruction of the core complex bound to gapped DNA.**

(A) Flow chart of cryo-EM image processing.

(B) Global FSC curves. The overall cryo-EM map resolution is 3.3 Å with FSC set at 0.143.

(C, D) Euler angle distribution (panel C) and final 3D reconstructed map colored according to local resolution (panel D).

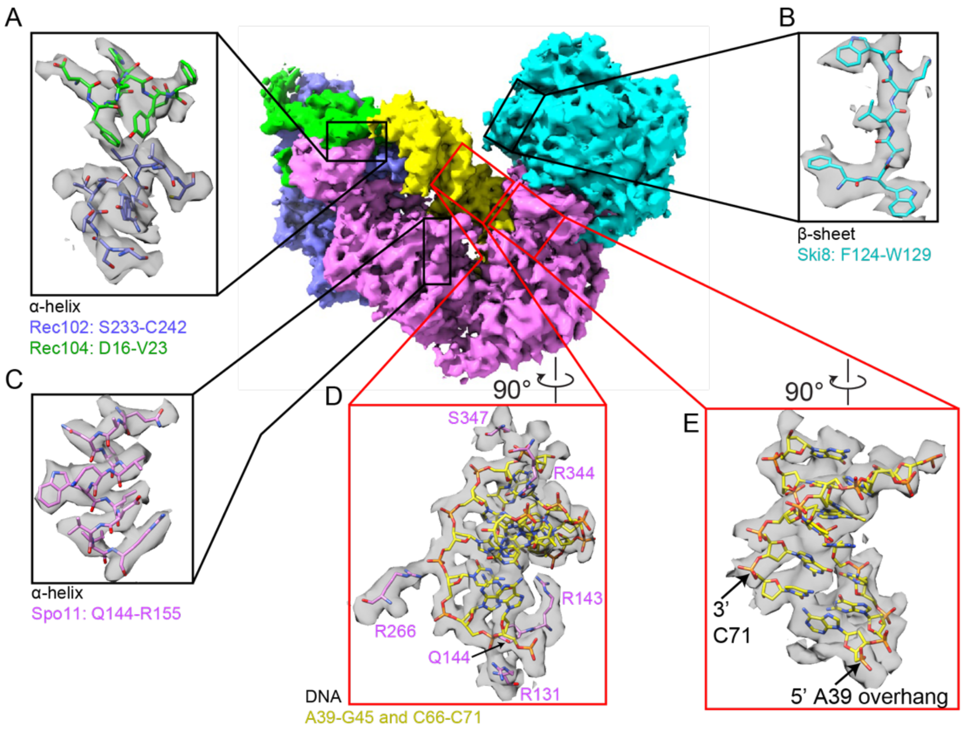


**Fig. S4. Examples of protein side chain and DNA base-sugar-phosphate identification in the cryo-EM structure of the core complex bound to gapped DNA.**

(A–E) The expanded boxes show examples of fitting of protein amino acid side chain (panels A–C) and DNA base-sugar-phosphate and interacting protein side chains (panels D and E) into the density map.

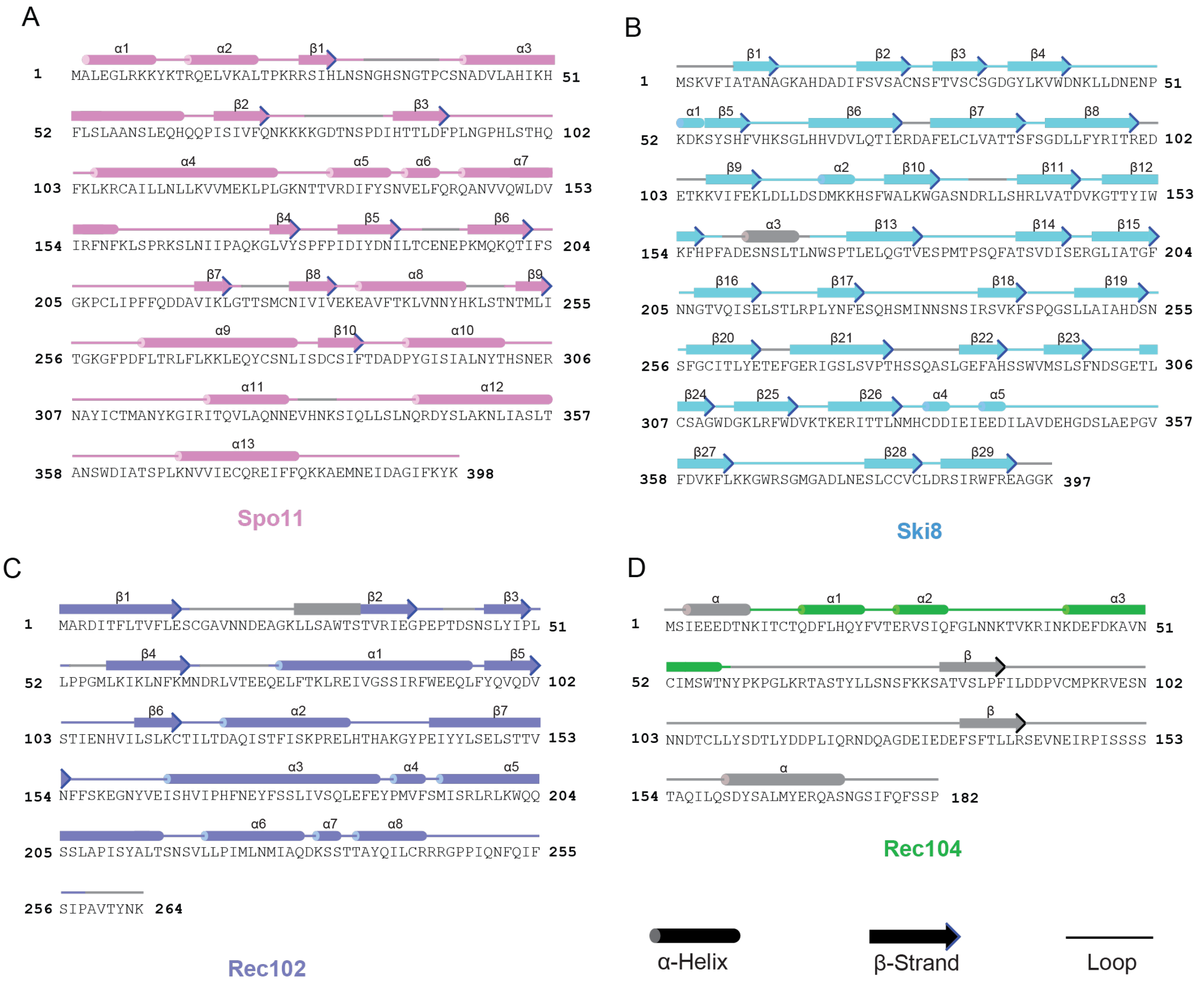


**Fig. S5. Protein secondary structures.**

(A–D) The secondary structures for Spo11 (panel A), Ski8 (panel B), Rec102 (panel C), and Rec104 (panel D) were determined from the cryo-EM structure of the core complex bound to gapped DNA. Gray-colored regions are not visible in the cryo-EM structure.

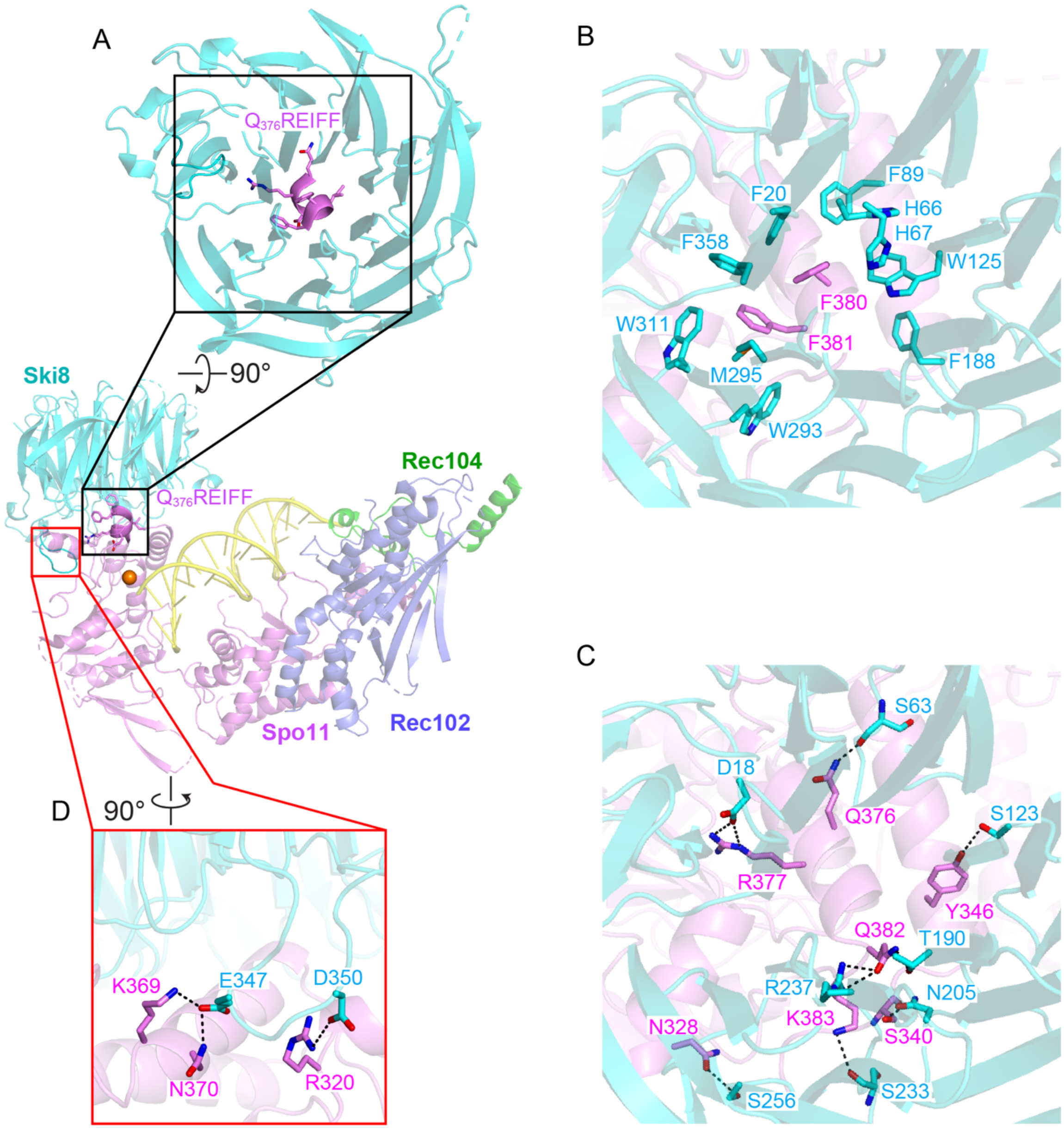


**Fig. S6. Interactions between Ski8 and Spo11.**

All images are from the cryo-EM structure of the core complex bound to gapped DNA.

(A) QREIFF motif from Spo11 (magenta) and the WD40 repeats from Ski8 (cyan).

(B) Hydrophobic interactions between Ski8 WD40 repeats and Spo11.

(C) Hydrogen bond interactions between Ski8 and Spo11.

(D) Additional hydrogen bonding interactions between Spo11 and the extended loop in Ski8.

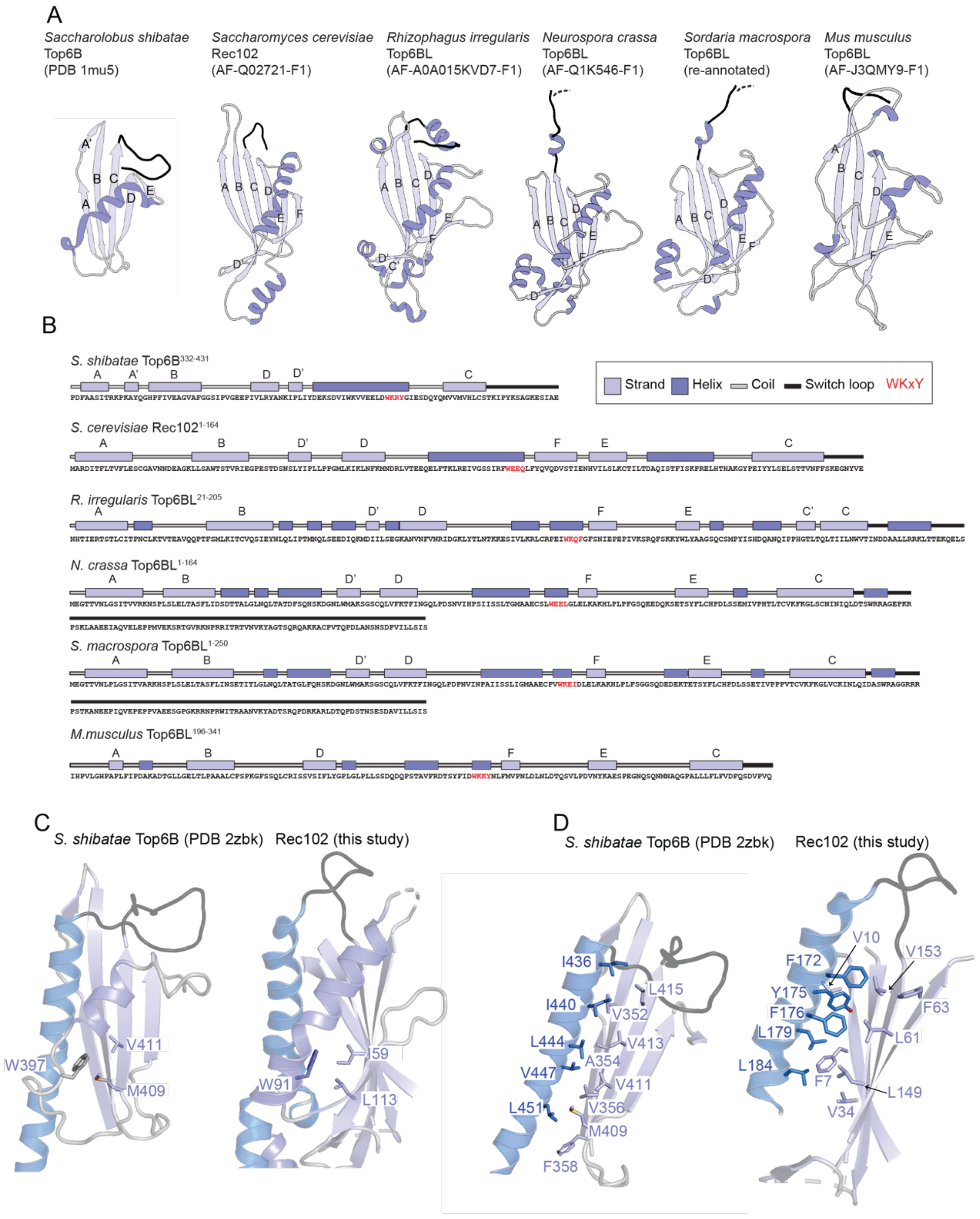


**Fig. S7. Sequence and structural conservation of Rec102 and Top6B.**

(A) Top6B and Rec102/Top6BL architectures. Strands, helices, and switch loops are colored as in **Fig. 3B** and strands are labeled as in **Fig. 3A**. All structures show truncated proteins, starting at the first β strand (strand A) and ending with the switch loop before the start of the stalk. The switch loop is truncated for the *N. crassa* and *S. macrospora* proteins for clarity.

(B) Secondary structure and sequence information for models shown in (A). The conserved WKxY motif is highlighted in red and β strands are labeled as in (A).

(C) Packing for *S. shibatae* Top6B (PDB: 2zbk) in comparison to Rec102 for the W in the WKxY motif. Ribbons and sticks are colored as in **Fig. 3B**.
(D) Detail view of the hydrophobic interface between the amphipathic stalk and the β strands; related residues in Top6B and Rec102 are indicated.

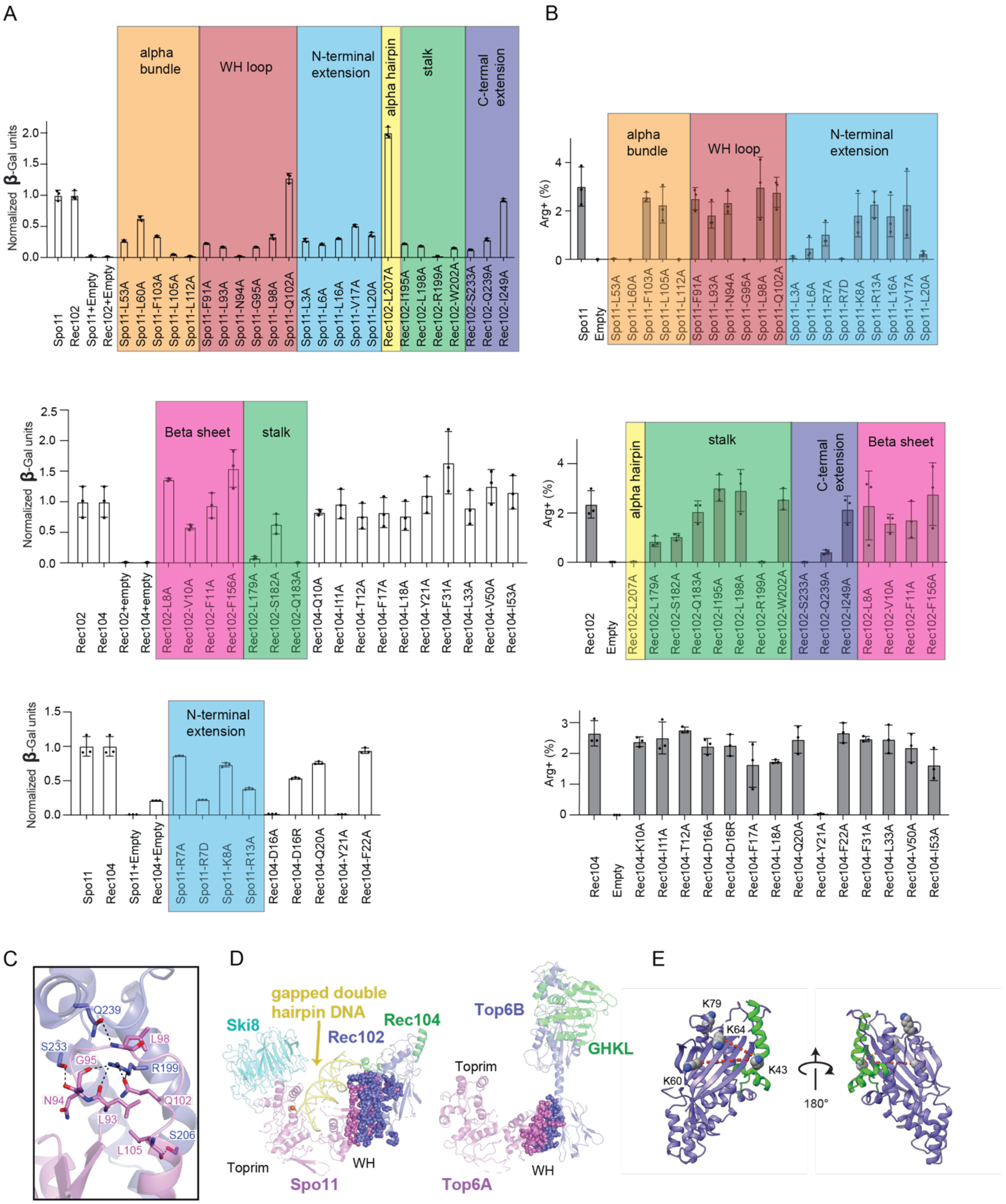


**Fig. S8. Functional analyses of protein-protein interfaces within the core complex.**
(A) Quantitative β-galactosidase assays to measure Y2H interactions of mutants of GAL4AD-Spo11 and LexA-Rec102 (top), LexA-Rec102 and GAL4AD-Rec104 (middle) and GAL4AD-

Spo11 and Rec104-LexA (bottom). The interaction of Rec102 with Rec104 was measured in vegetative conditions and the rest were measured in meiotic conditions.
(B) Heteroallele recombination assays with different Spo11 (top), Rec102 (middle) and Rec104 (bottom) mutant proteins. The graphs show the frequency of Arg+ prototrophs generated by recombination between *arg4-bgl* and *arg4-nsp* alleles[54]. In panels A and B, bars show mean ± SD of three replicates; points show individual measurements.
(C) Hydrogen bonding interactions between Spo11 and Rec102.
(D) Comparison of interfacial areas for Spo11 with Rec102 (1,680 Å$^2$, complex with gapped DNA) vs. Top6A with Top6B from *M. mazei* Topo VI (958 Å$^2$, PDB: 2q2e). Residues involved in the interfaces are shown as spheres.
(E) Positions of lysines in Rec102 and Rec104 that are readily crosslinked. Red dashed lines connect the $\alpha$ carbons of lysines that were crosslinked in[14].

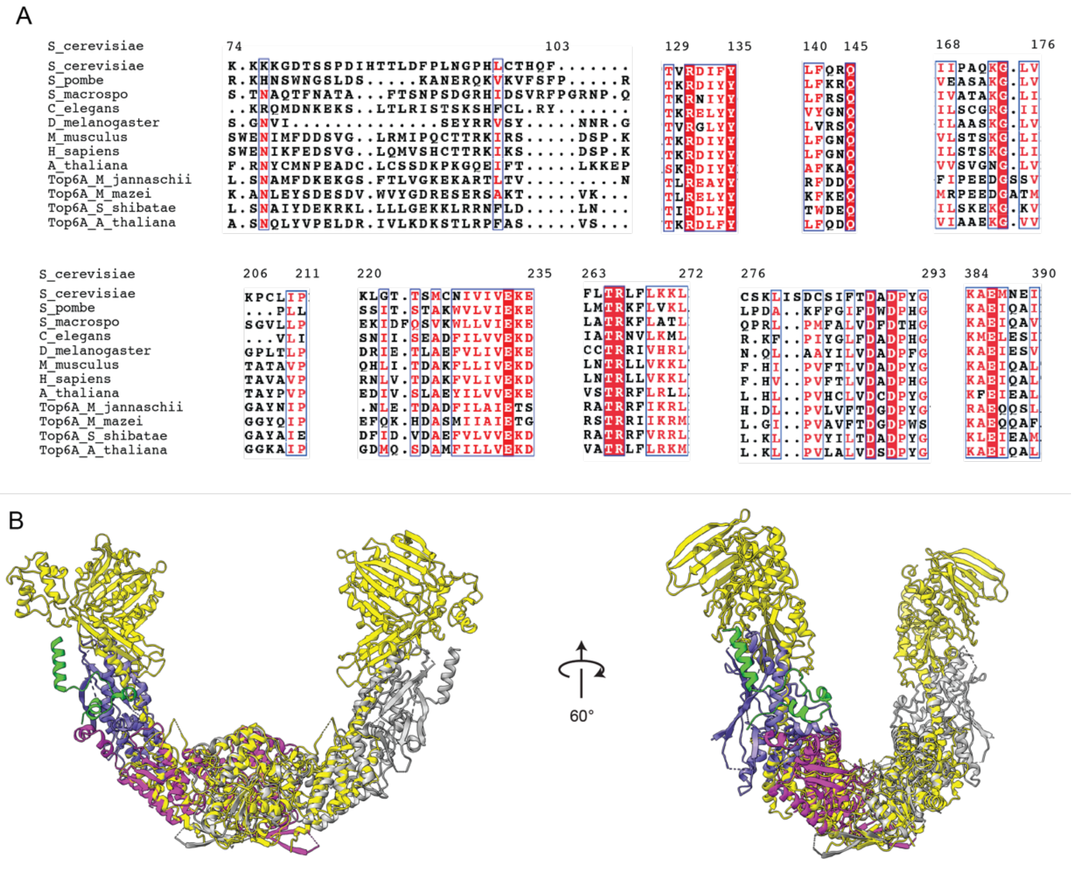


**Fig. S9. Spo11 sequence conservation.**
(A) Multiple sequence alignment of Spo11 and Top6A orthologs at protein-protein and protein-DNA contact sites.
(B) Structural alignment of the hypothetical model of a Spo11 core complex pre-DSB dimer and the crystal structure of the dimeric *S. shibatae* Topo VI holoenzyme (yellow). One copy of the Spo11 core complex is colored as **Fig. 1B**, and the other copy is colored in gray. Ski8 is not shown for simplicity.

### SUPPLEMENTAL TABLES

#### Table S1. Cryo-EM data collection, processing, and validation statistics

| | Spo11 core complex with: | |
|---|---|---|
| | **Hairpin DNA** | **Gapped DNA** |
| Data collection | | |
| Microscope | Titan Krios | Titan Krios |
| Detector | Gatan K3 | Gatan K3 |
| Automation software | SerialEM | SerialEM |
| Nominal magnification | 22,500 | 22,500 |
| Voltage (kV) | 300 kV | 300 kV |
| Total dose ($e^-$/Å$^2$) | 53.00 | 53.00 |
| Dose rate ($e^-$/pixel/s) | 20 | 20 |
| Number of frames collected | 40 | 40 |
| Defocus range (μm) | -1.0 to -2.5 | -1.0 to -2.5 |
| Pixel size (Å) | 1.064 | 1.064 |
| Collected Micrographs | 3,258 | 5,521 |
| Selected Micrographs | 2,388 | 5,221 |
| Reconstruction | | |
| Initially autopicked particles | 1,829,931 | 7,804,373 |
| Particles used for classification | 1,284,897 | 6,930,069 |
| Particles in the final map | 128,787 | 548,674 |
| Symmetry | C1 | C1 |
| Resolution | | |
| FSC 0.5 (unmasked/masked, Å) | 4.1/3.9 | 3.7/3.6 |
| FSC 0.143 (unmasked/masked, Å) | 3.7/3.6 | 3.3/3.3 |
| Map sharpening B factor (Å$^2$) | -158.0 | -155.8 |
| Model composition | | |
| Protein residues | 1001 | 993 |
| Nucleotide | 43 | 31 |
| Nonhydrogen atoms | 8895 | 8569 |
| Validation | | |
| MolProbity | 2.80 | 2.84 |
| Clash score | 20 | 18 |
| Map Correlation Coefficient | 0.74 | 0.71 |
| R.m.s. deviations | | |
| Bond lengths (Å) | 0.005 | 0.017 |
| Bond angles (°) | 0.979 | 1.250 |
| Ramachandran plots | | |
| Favored (%) | 88.11 | 88.88 |
| Allowed (%) | 10.95 | 10.81 |
| Outliers (%) | 0.94 | 0.31 |
| Rotamer outliers (%) | 3.21 | 4.61 |

**Table S2. Yeast strains**

| Strain | Genotype |
|---|---|
| SKY661 | *MATa, ho::LYS2, lys2, leu2::hisG, trp1::hisG, ndt80::kanMX, LexA(op)-LacZ::URA3* |
| SKY662 | *MATα, ho::LYS2, lys2, leu2::hisG, trp1::hisG, ndt80::kanMX, LexA(op)-LacZ::URA3* |
| SKY1311 | *MATα, ho::LYS2, lys2, ura3, leu2::hisG, trp1::hisG, arg4-nsp, rec102∆::URA3* |
| SKY1312 | *MATa, ho::LYS2, lys2, ura3, leu2::hisG, trp1::hisG, arg4-bgl, rec102∆::URA3* |
| SKY969 | *MATα, ho::LYS2, lys2, leu2::hisG, trp1::hisG, arg4-bgl, spo11Δ::hisG-URA3-hisG* |
| SKY970 | *MATa, ho::LYS2, lys2, leu2::hisG, trp1::hisG, arg4-nsp, spo11Δ::hisG-URA3-hisG* |
| SKY7404 | *MATa, ho::LYS2, lys2, leu2::hisG, trp1::hisG, arg4-bgl, rec104Δ::KanMX* |
| SKY7405 | *MATα,ho::LYS2, lys2, leu2::hisG, trp1::hisG, arg4-nsp, rec104Δ::KanMX* |

### Table S3. Plasmids

| Plasmid | Description | Reference |
|---|---|---|
| pCCB586 | *SPO11* from *S. cerevisiae* (SK1 strain) in pFastBac1 | Claeys Bouuaert et al., 2021 |
| pCCB587 | *SKI8* from *S. cerevisiae* (SK1 strain) in pFastBac1 | Claeys Bouuaert et al., 2021 |
| pCCB588 | *REC102* from *S. cerevisiae* (SK1 strain) in pFastBac1 | Claeys Bouuaert et al., 2021 |
| pCCB589 | *REC104* from *S. cerevisiae* (SK1 strain) in pFastBac1 | Claeys Bouuaert et al., 2021 |
| pSK275 | LexA empty Y2H vector | Arora et al., 2004 |
| pSK276 | Gal4AD empty Y2H vector | Arora et al., 2004 |
| pSK282 | LexA-Rec102 Y2H vector (pCA1-Rec102) | Maleki et al., 2007 |
| pSK293 | Rec104-LexA Y2H vector | Arora et al., 2004 |
| pSK305 | Gal4AD-Spo11 Y2H vector | Arora et al., 2004 |
| pSK310 | Gal4AD-Rec104 Y2H vector | Arora et al., 2004 |
| pZZy1 | pSK305 with *spo11-L53A* | This study |
| pZZy2 | pSK305 with *spo11-L60A* | This study |
| pZZy3 | pSK305 with *spo11-F91A* | This study |
| pZZy4 | pSK305 with *spo11-L105A* | This study |
| pZZy5 | pSK305 with *spo11-L112A* | This study |
| pZZy6 | pSK305 with *spo11-L93A* | This study |
| pZZy7 | pSK305 with *spo11-N94A* | This study |
| pZZy8 | pSK305 with *spo11-G95A* | This study |
| pZZy9 | pSK305 with *spo11-L98A* | This study |
| pZZy10 | pSK305 with *spo11-F103A* | This study |
| pZZy11 | pSK305 with *spo11-L3A* | This study |
| pZZy12 | pSK305 with *spo11-R6A* | This study |
| pZZy13 | pSK305 with *spo11-R7A* | This study |
| pZZy14 | pSK305 with *spo11-R7D* | This study |
| pZZy15 | pSK305 with *spo11-L16A* | This study |
| pZZy16 | pSK305 with *spo11-V17A* | This study |
| pZZy17 | pSK305 with *spo11-L20A* | This study |
| pZZy18 | pSK282 with *rec102-L207A* | This study |
| pZZy19 | pSK282 with *rec102-I195A* | This study |
| pZZy20 | pSK282 with *rec102-L198A* | This study |
| pZZy21 | pSK282 with *rec102-R199A* | This study |

| | | |
|---|---|---|
| pZZy22 | pSK282 with *rec102-W202A* | This study |
| pZZy23 | pSK282 with *rec102-S233A* | This study |
| pZZy24 | pSK282 with *rec102-I249A* | This study |
| pZZy25 | pSK282 with *rec102-L8A* | This study |
| pZZy26 | pSK282 with *rec102-V10A* | This study |
| pZZy27 | pSK282 with *rec102-L179A* | This study |
| pZZy28 | pSK282 with *rec102-S182A* | This study |
| pZZy29 | pSK282 with *rec102-F11A* | This study |
| pZZy30 | pSK282 with *rec102-F156A* | This study |
| pZZy31 | pSK310 with *rec104-I11A* | This study |
| pZZy32 | pSK310 with *rec104-T12A* | This study |
| pZZy33 | pSK310 with *rec104-F17A* | This study |
| pZZy34 | pSK310 with *rec104-L18A* | This study |
| pZZy35 | pSK310 with *rec104-Y21A* | This study |
| pZZy36 | pSK310 with *rec104-F31A* | This study |
| pZZy37 | pSK310 with *rec104-L33A* | This study |
| pZZy38 | pSK310 with *rec104-V50A* | This study |
| pZZy39 | pSK310 with *rec104-I53A* | This study |
| pZZy40 | pSK293 with *rec104-D16A* | This study |
| pZZy41 | pSK293 with *rec104-D16R* | This study |
| pZZy42 | pSK293 with *rec104-Y21A* | This study |
| pZZy43 | pSK305 with *spo11-R13A* | This study |
| pZZy44 | pSK305 with *spo11-K8A* | This study |
| pZZy45 | pSK282 with *rec102-Q183A* | This study |
| pZZy46 | pSK305 with *spo11-Q012A* | This study |
| pZZy47 | pSK282 with *rec102-Q239A* | This study |
| pZZy48 | pSK293 with *rec104-F22A* | This study |
| pZZy49 | pSK293 with *rec104-Q20A* | This study |
| pZZy50 | pSK310 with *rec104-K10A* | This study |
